# Type-I IFNs induce GBPs and lysosomal defense in hepatocytes to control malaria

**DOI:** 10.1101/2024.10.22.619707

**Authors:** Camila Marques-da-Silva, Clyde Schmidt-Silva, Carson Bowers, Essel Charles-Chess, Justine C. Shiau, Eui-Soon Park, Zhongyu Yuan, Bae-Hoon Kim, Dennis E. Kyle, John T. Harty, John D. MacMicking, Samarchith P. Kurup

**Affiliations:** Department of Cellular Biology, University of Georgia, Athens, GA, USA; Center for Tropical and Emerging Global Diseases, University of Georgia, Athens, GA; Department of Infectious Diseases, University of Georgia, Athens, GA; Howard Hughes Medical Institute, Yale University School of Medicine, New Haven, CT; Yale Systems Biology Institute, West Haven, CT; Department of Microbial Pathogenesis, Yale University School of Medicine, New Haven, CT; Department of Immunobiology, Yale University School of Medicine, New Haven, CT; Department of Pathology, University of Iowa, Iowa City, IA; Interdisciplinary Graduate Program in Immunology, University of Iowa, Iowa City, IA

## Abstract

*Plasmodium* parasites undergo development and replication within the hepatocytes before infecting the erythrocytes and initiating clinical malaria. Although type-I interferons (IFNs) are known to hinder *Plasmodium* infection within the liver, the underlying mechanisms remain unclear. Here, we describe two IFN-I-driven hepatocyte antimicrobial programs controlling liver-stage malaria. First, oxidative defense by NADPH oxidases 2 and 4 triggers a pathway of lysosomal fusion with the parasitophorous vacuole (PV) to help clear *Plasmodium*. Second, guanylate-binding protein (GBP) 1 disruption of the PV activates caspase-1 inflammasome, inducing pyroptosis to remove the infected host cells. Remarkably, both human and mouse hepatocytes enlist these cell-autonomous immune programs to eliminate *Plasmodium*; their pharmacologic or genetic inhibition led to profound malarial susceptibility, and are essential *in vivo*. In addition to identifying the IFN-I-mediated cell-autonomous immune circuits controlling *Plasmodium* infection in the hepatocytes, this study extends our understanding of how non-immune cells are integral to protective immunity against malaria.

## Introduction

Malaria remains a significant global health crisis affecting over half of the world’s population (WHO, 2023). The *Plasmodiu*m parasite responsible is inoculated by mosquitoes as sporozoites (Spz) into the skin of mammalian hosts where they undergo development and replication within hepatocytes (liver-stage malaria) before infecting erythrocytes and initiating a potentially lethal ‘blood-stage’ of disease. The Spz develops within a parasitophorous vacuole (PV) enveloped by the PV membrane (PVM) inside host cells. This intracellular niche is believed to isolate and protect the parasite within the infected hepatocytes (Bano et al., 2007; Nyboer et al., 2018).

Hepatocytes constitute over 90% of the cellular biomass of the liver and have traditionally been viewed as immunologically inert bystanders in the context of hepatotropic infections (Kubes and Jenne, 2018). However, recent studies have highlighted their role in the innate and adaptive immune control of such infections, including malaria (Bogdanos et al., 2013; Bolen et al., 2014; Crispe, 2016; Kurup et al., 2019a; Lefebvre et al., 2021; Marques-da-Silva et al., 2023; Tretina et al., 2019).

Type-I IFNs drive protective responses against liver-stage malaria, with hepatocytes themselves producing IFN-β that acts locally to reduce overall *Plasmodium* burdens in the liver (Gazzinelli et al., 2014; He et al., 2020; Liehl et al., 2014; Marques-da-Silva et al., 2022; Marques-da-Silva *et al*., 2023; Marques-da-Silva et al., 2024b; Miller et al., 2014; Nussler et al., 1993). Human and murine primary hepatocyte cultures, as well as whole livers of mice, exhibit transcriptional signatures associated with type-I IFN stimulation following *Plasmodium* infection (He *et al*., 2020; Liehl *et al*., 2014; Marques-da-Silva *et al*., 2022; Marques-da-Silva *et al*., 2023; Miller *et al*., 2014). Treatment of *P. berghei* ANKA (*Pb*)*, P. falciparum* (*Pf*), or *P. vivax* (*Pv*)*-*infected murine or human hepatocytes with sufficient IFN-α or IFN-β limited these infections (Mancio-Silva et al., 2022; Marques-da-Silva *et al*., 2022). Conditional deletion of the IFNαα/β receptor (IFNAR) specifically in the infected hepatocytes using cre-recombinase-expressing *Pb* resulted in higher overall parasite burdens in the liver and ablated the protection conferred by exogenous IFN-I treatment (Marques-da-Silva *et al*., 2022). Thus, direct type-I IFN signaling within infected hepatocytes appears essential to control *Plasmodium* infection in the liver. How type-I IFNs exert such control, however, is currently unclear.

It is known that hepatocytes respond to IFNs by activating a variety of interferon- stimulated genes (ISGs), including NADPH oxidases (NOXs), which generate reactive oxygen species (ROS) (Bolen *et al*., 2014; Dandri et al., 2021; Kumova and Carey, 2020; MacMicking, 2012; Tretina *et al*., 2019; Watanabe et al., 2003; Xu et al., 2021). ROS is considered most effective against vesiculated pathogens through the induction of the LC3-associated phagocytosis (LAP) pathway (Heckmann and Green, 2019; Paiva and Bozza, 2014). LAP is characterized by the presence of a single-membraned phagosome (distinguishing it from autophagy-derived double-membrane structures), NOX-mediated ROS production, and localization of the regulator RUBICON (RUN domain protein interacting with Beclin-1 and cysteine-rich, RUBCN) to the phagosome (Martinez et al., 2015; Martinez-Sanchez et al., 2018). ROS stimulates lipidation of the microtubule- associated protein 1 light chain 3 (LC3) to generate LC3-II, which binds to the phagosome, marking the latter for lysosomal fusion (Heckmann and Green, 2019; Wang et al., 2022). Lysosomal fusion leads to the acidification and destruction of the phagosomal contents (Heckmann and Green, 2019; Martinez-Sanchez et al., 2018).

Lysosomes have been observed to sporadically associate with the PVM of *Pb* and *Pv* in hepatocytes (Boonhok et al., 2016; Niklaus et al., 2019; Prado et al., 2015) where they may facilitate *Plasmodium* control (Petersen et al., 2017).

IFNs also induce the expression of guanylate-binding proteins (GBPs) (Kim et al., 2016; Tretina *et al*., 2019) which bind to host and pathogen membranes to mechano- enzymatically disrupt them, often exposing the contents of the vacuole and the pathogen to the cytosol (Bass et al., 2023; Kim et al., 2012; Kravets et al., 2016; Weismehl et al., 2024; Zhu et al., 2024). GBPs also seed the formation of large supramolecular complexes that promote the activation of allosterically weaker enzymes such as caspases, in some cases directly on the surface of targeted microbes (Fisch et al., 2020; Santos et al., 2018; Shenoy et al., 2012; Wandel et al., 2020; Zhu *et al*., 2024). Various GBPs have been shown to protect host cells from intracellular pathogens such as *Chlamydia, Fransicella, Legionella, Mycobacteria, Salmonella, Shigella,* or *Toxoplasma* (Feeley et al., 2017; Finethy et al., 2015; Haldar et al., 2014; Kim et al., 2011; Selleck et al., 2013; Tretina *et al*., 2019; Wandel *et al*., 2020). Some of the above protection is exerted through canonical caspase-1 or non-canonical caspase-4/11-driven inflammasome activation, including downstream of the innate immune sensor, absent in melanoma (AIM)-2 (Man et al., 2016; Meunier et al., 2015).

*Plasmodium* double-strand (ds) DNA exposed to the host cell cytoplasm induces AIM2 inflammasome-mediated caspase-1 activation and pyroptosis, resulting in the release of *Plasmodium* antigens to the extracellular milieu and the generation of *Plasmodium*-specific adaptive immune responses (Kurup *et al*., 2019a; Marques-da-Silva *et al*., 2023; Marques-da-Silva et al., 2024a). Similarly, *Plasmodium* RNA released into the hepatocyte cytosol generate additional IFN-I, as well as the innate and adaptive control of malaria (Liehl *et al*., 2014; Miller *et al*., 2014). Although type-I IFN-driven lysis of *Plasmodium* and its PV are potentially responsible for triggering such pathways, how they do so has remained unknown (**Fig 1A**). This is what we set out to investigate.

**Figure 1.**
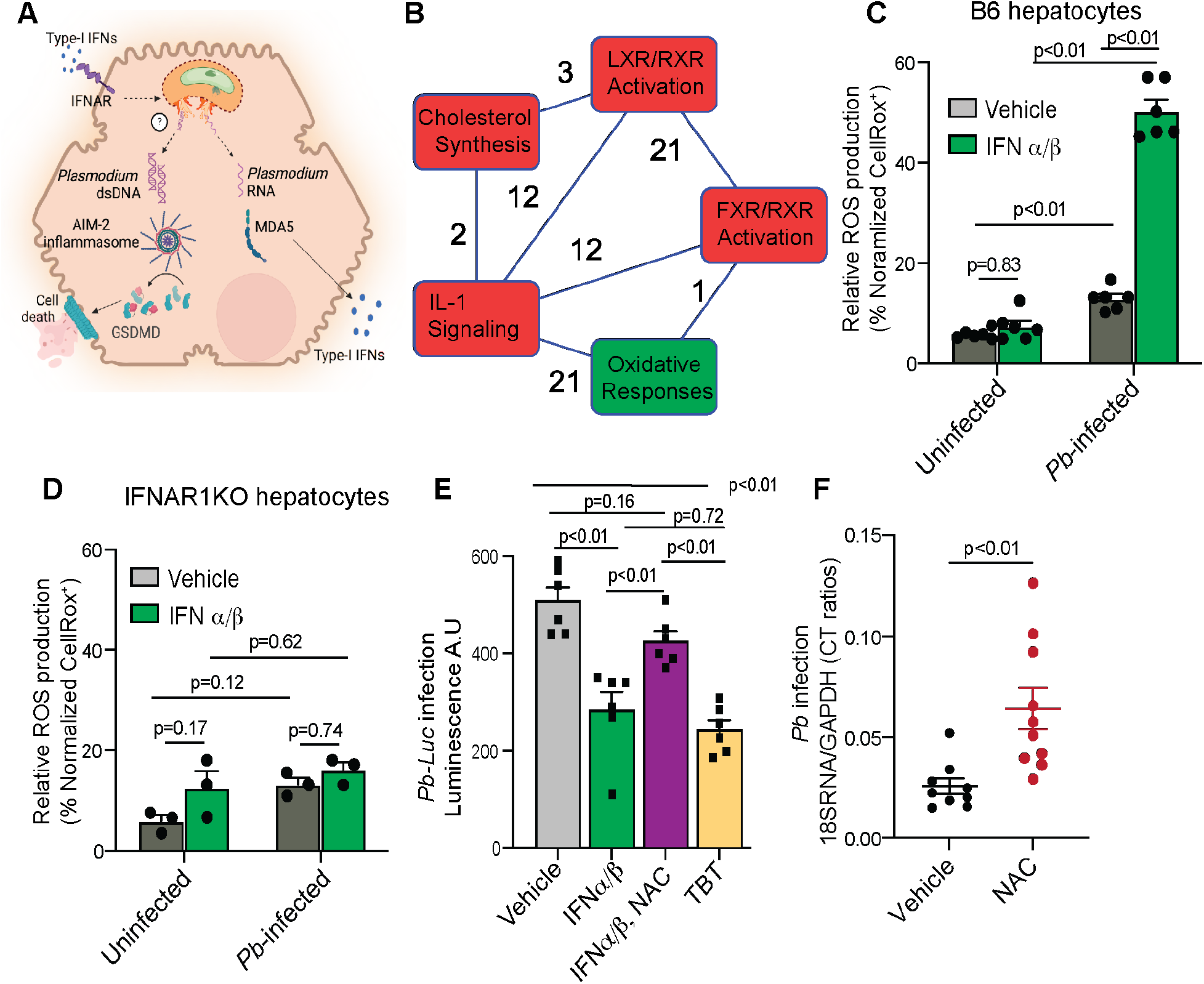
Type-I IFNs induce ROS-mediated control of *P. berghei* infection in hepatocytes *in vitro* and *in vivo*. (A) Current model for type-I IFN response in *Plasmodium*-infected hepatocytes. Type-I IFN stimulation in *Plasmodium*-infected hepatocytes potentially triggers, through some unknown mechanisms, the lysis of the parasite and the PVM to aid the release of parasite RNA or dsDNA into the host cell cytoplasm, driving additional type-I IFN production or pyroptosis respectively. (B) Primary canonical biochemical pathways perturbed in *Pb-GFP*-infected primary B6 mouse hepatocytes determined by gene-set enrichment analysis of transcripts through the gene ontology algorithms of Ingenuity Pathway Analysis (IPA), 36h p.i. The connecting lines and the corresponding numbers indicate the number of individual overlapping genes. Source data: NCBI Gene Expression Omnibus GSE186023. (C-D) ROS in *Pb*-infected B6 (B) or IFNARKO (C) hepatocyte cultures determined 24h p.i., following treatment with vehicle (PBS) or IFNα/β, 6h p.i. Combined data presented as mean + s.e.m with each data point obtained from a separate replicate experiment, analyzed using 2-way ANOVA with Tukey’s correction to yield the indicated p-values. (E) Relative infection loads in primary B6 hepatocyte cultures infected with *Pb-Luc* quantified by total luminescence signal at 32h p.i., with the indicated treatments. Combined data presented as mean + s.e.m with each data point obtained from a separate experiment, analyzed using 1-way ANOVA with Tukey’s correction to yield the indicated p-values. (F) Scatter plot indicating relative liver parasite burdens in B6 mice inoculated with *Pb* and treated with vehicle control or NAC from −24hp.i., determined by qRT-PCR at 40h p.i. Combined data from replicate experiments with each data point representing a mouse, presented as mean + s.e.m and analyzed using unpaired t-test to obtain the indicated p-values.

## Results

### Autocrine or paracrine control *Plasmodium* infection by IFN-I

Type-I IFNs generated by infected hepatocytes is believed to facilitate *Plasmodium* control in the liver (Liehl *et al*., 2014; Marques-da-Silva *et al*., 2022); increasing inoculums of *Plasmodium* Spz should therefore produce enough type-I IFNs to autocrinally diminish parasite loads in culture. To test this idea, we inoculated primary C57BL/6 (B6) mouse hepatocyte cultures with increasing doses of *Pb* Spz. Our choice of primary rather than immortalized hepatic cell lines is based on the latter having altered immune, signaling, cell-death, and metabolic pathways (Fridman and Tainsky, 2008; Garcia et al., 2006; Liu et al., 2018; Wang et al., 2011).

We found IFN-α levels in the supernatants significantly increased with parasite inoculum (**Fig S1A**) and coincided with curtailed infection in B6 hepatocytes at higher inoculums (**Fig S1B**). Such curtailment was absent in IFNAR-deficient (IFNARKO) hepatocyte cultures (**Fig S1B**), confirming that type-I IFN signaling was responsible for the decline in frequencies of infected hepatocytes. Notably, the fraction of hepatocytes clearing *Pb* infection in mouse livers is also increased with higher Spz inoculums (Afriat et al., 2022). Our findings indicate that cultured primary hepatocytes inoculated with *Plasmodium* generate type-I IFNs in a dose-dependent manner and induce downstream IFN-I signaling to control infection *in vitro*.

### IFNα/β-induced hepatocyte ROS is essential for *Plasmodium* control *in vitro* and *in vivo*

Analysis of our published transcriptional data (GSE186023 (Marques-da-Silva *et al*., 2022)) for genes with >2-fold expression changes indicated that the oxidative stress response pathway was particularly enriched in hepatocyte cultures following *Pb* infection (**Fig 1B**). Indeed, *Pb*-infected hepatocytes in culture generated ROS, which was further augmented by supplementation with exogenous recombinant IFN-α and IFN-β (IFNα/β, **Fig 1C**). Such induction of ROS was absent in IFNAR1KO hepatocytes (**Fig 1D**).

To test if ROS impacted *Plasmodium* control, we infected hepatocyte cultures with a strain of *Pb* expressing luciferase (*Pb-Luc*) and quantified the total luminescence signal. While exogenous IFNα/β induced lower parasite burdens in the *Pb-Luc*-inoculated cultures, this was reversed by treatment with the ROS scavenger, N-acetyl cysteine (NAC, **Figs S2A, 1E**). Conversely, the treatment with an inducer of intracellular ROS, Tributyltin (TBT) facilitated parasite killing, akin to IFNα/β treatment (**Fig 1E**). These *in-vitro* findings were corroborated by the presence of significantly higher liver parasite burdens in *Pb*-infected B6 mice pre-treated with NAC (**Fig 1F**). Intriguingly, the difference in parasite loads following NAC treatment was more conspicuous in the earlier stages of the infection, before schizogony (**Figs S2B, S2C**). Taken together, our data indicated that IFN-I-induced ROS is a key driver of *Pb* control in hepatocytes.

Two major sources of ROS in hepatocytes are NOX2 and NOX4. Both were significantly upregulated in B6 hepatocyte cultures as well as whole mouse livers following *Pb* infection (**Fig 2A-E**). Moreover, they also localized to *Pb* in hepatocytes, the frequencies of which were enhanced by the addition of exogenous IFNα/β (**Fig 2F-I**). To determine the functional impact of NOX2 or NOX4 on *Plasmodium* control, we employed their specific therapeutic inhibitors, GSK2795039 (Hirano et al., 2015) or GLX351322 (Anvari et al., 2015), respectively. Both the drugs impeded the generation of ROS in *Pb*-infected hepatocyte cultures in a dose-dependent manner (**Fig S2D**) and reversed IFN-I-mediated control of *Pb* infection (**Fig 2J**). Of note, combining GSK2795039 and GLX351322 appeared to have a cumulatively higher impact on limiting *Pb* infection, suggesting that NOX2 and NOX4 may have non-redundant roles in ROS production in *Pb*-infected hepatocytes. GSK2795039 and GLX351322 also hindered IFN-I-mediated control of *Pf* infection in primary human hepatocytes (**Fig 2K**). These *in vitro* findings were reinforced by the observation of higher liver parasite burdens in *Pb*-infected B6 mice treated with NOX2 and NOX4 inhibitors (**Fig 2L**). Although *Pb* infection is acutely lethal in B6 mice, adequate dosing with IFNα/β clears it from the liver, yielding 100% survival of the infected mice (Marques-da-Silva *et al*., 2022). Blocking NOX2 and NOX4 reversed this IFN-I-driven protection, resulting in 100% mortality (**Fig 2M**).

**Figure 2.**
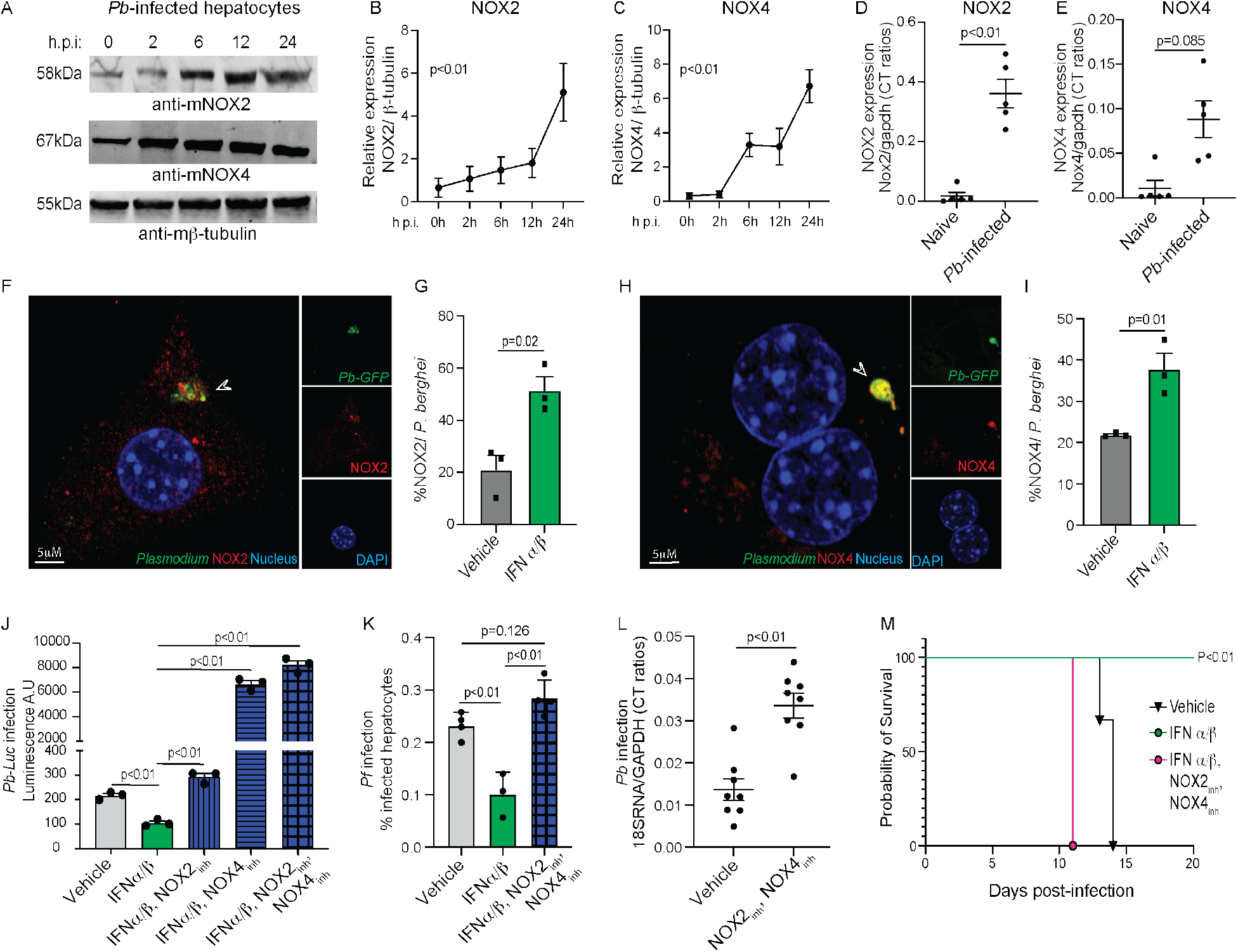
NOX2 and NOX4 induced by type-I IFNs in *Plasmodium*-infected hepatocytes essential for controlling liver-stage malaria. (A) Immunoblots depicting the expression kinetics of NOX2 or NOX4 in primary mouse (B6) hepatocyte cultures inoculated with *Pb*. β-tubulin served as loading controls, data representative of 4 replicate experiments. (B-C) Kinetics of NOX2 (B) or NOX4 (C) expression quantified from (A) using relative band densities using β-tubulin as loading control. Data from 4 replicate experiments presented as mean + s.e.m, analyzed using ANOVA with Geisser-Greenhouse correction to obtain the indicated p-value. (D-E) Scatter plots indicating NOX2 (D) or NOX4 (E) transcript levels determined by qRT- PCR in hepatocytes isolated from *Pb-*infected B6 mice, 12h p.i. Data combined from two experiments presented as mean + s.e.m with each data point representing a mouse, analyzed using unpaired t-test to obtain the indicated p-values. (F-I) Pseudo-colored confocal images depicting the subcellular localization of NOX2 (F) or NOX4 (H) in primary B6 hepatocytes infected with *Pb-GFP*, 24h p.i. Image obtained from one of >3 replicates, arrows indicate *Pb* EEFs. Bar graphs depict the frequencies of NOX2 (G) or NOX4 (I) co-localization with *Pb-GFP*, 26h p.i., in primary mouse (B6) hepatocytes treated with vehicle or IFNα/β at 24h p.i. Each data point represents a separate experiment where >50 infected hepatocytes were analyzed, presented as mean + s.e.m, and analyzed using unpaired t-test to obtain the indicated p-values. (J) Relative infection loads in primary mouse (B6) hepatocyte cultures inoculated with *Pb- Luc* with the indicated treatments from 6h p.i., quantified by total luminescence signal at 32h p.i. (K) Relative infection loads in primary human hepatocyte cultures inoculated with *P. falciparum* and with the indicated treatments, quantified using high-content imaging, 4d p.i. J,K: Combined data presented as mean + s.e.m, with each point representing a separate experiment, analyzed using 1-way ANOVA with Dunnets (J) or Tukey’s (K) correction to obtain the indicated p-values. (L) Scatter plots indicating relative liver parasite burdens in *Pb*-inoculated B6 mice treated with vehicle or NOX inhibitors, determined by qRT-PCR, 40h p.i. Combined data from three separate experiments with each data point indicating a mouse, presented as mean + s.e.m, analyzed using unpaired t-test to obtain the indicated p-values. (M) Survival kinetics of *Pb*-infected B6 mice treated as indicated. Representative data from 2 experiments with 4 mice/group, analyzed using Mantel-Cox test to yield the indicated p-values. Vehicle-treated mice eventually die >32d p.i.

### Type-I IFNs induce the LAP in *Plasmodium*-infected hepatocytes to protect against malaria

Because ROS can induce LAP against pathogens enclosed in a single membrane vacuole like *Plasmodium* (Heckmann and Green, 2019; Paiva and Bozza, 2014), we considered the possibility of IFN-I inducing LAP within infected hepatocytes. Indeed, RUBCN, which delineates the initiation of LAP, was rapidly induced in *Pb*-infected hepatocytes (**Fig S2E, S2F**) and directly localized to *Pb* (**Fig S2G**). The frequency of such localization was significantly enhanced by exogenous IFNα/β (**Fig S2H**). Similarly, activation of LC3, characterized by the transformation of the 16kDa LC3-I to 14kDa LC3- II that binds to single-membraned LAP structures (**Fig S3A**), was evident in the *Pb*- infected hepatocytes (**Fig 3A**). Although primary murine hepatocytes express high baseline levels of LC3-II (Chen et al., 2014; Karim et al., 2007), we observed a temporally distinct increase in LC3-II levels in *Pb*-infected B6 hepatocyte cultures, peaking at approximately 12h p.i. (hours post-infection, **Fig 3B**). These LC3-II levels were further enhanced by exogenous IFNα/β treatment, which was reversed by inhibiting ROS or NOX4 (**Fig 3C, 3D**).

**Figure 3.**
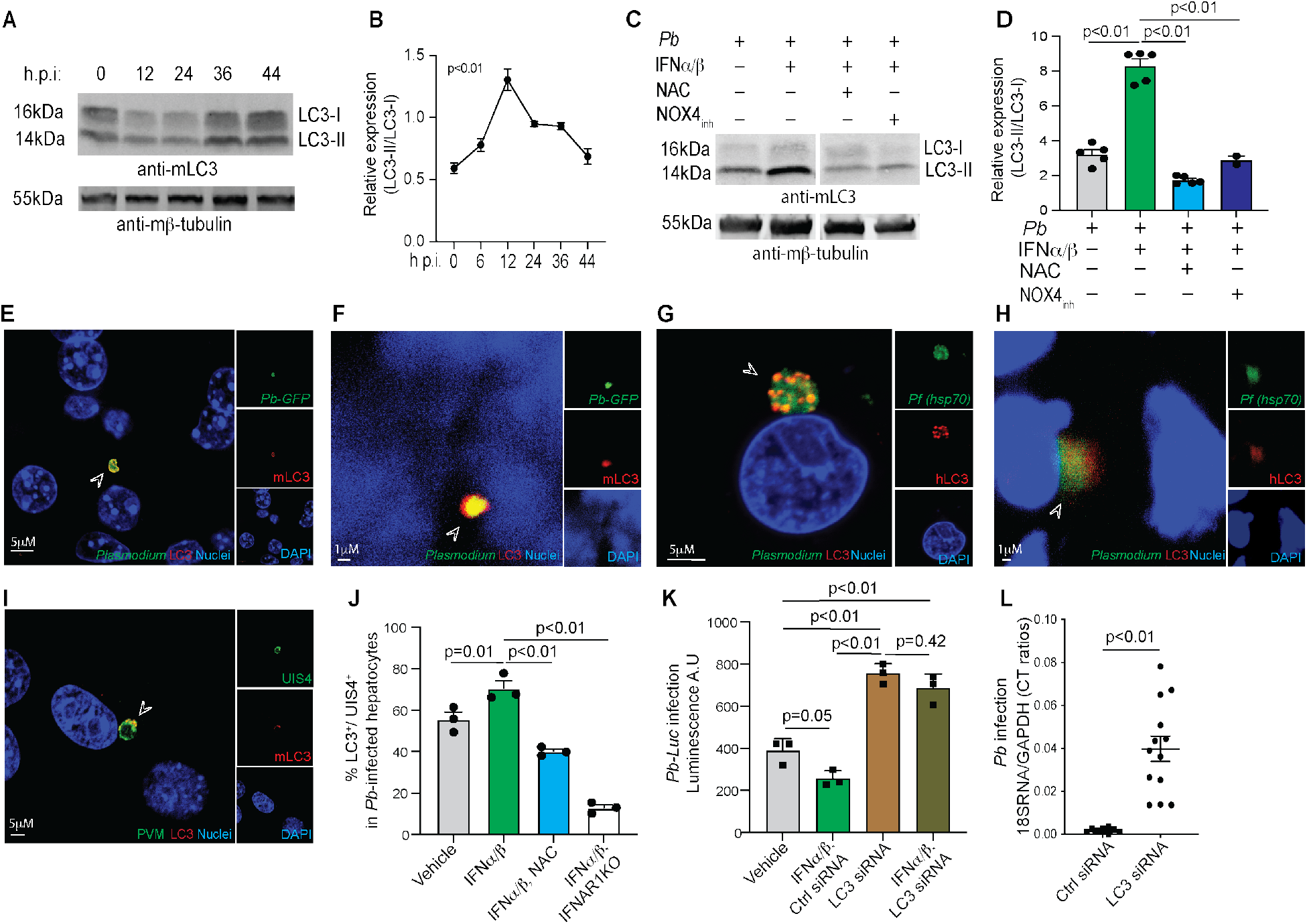
IFN-I induced LC3 activation and localization to *Plasmodium* in hepatocytes limits liver-stage malaria. (A) Immunoblots depicting LC3 expression in primary B6 mouse hepatocyte cultures co- incubated with *Pb* for the indicated time-frames. β-tubulin served as loading controls, data representative of 4 replicate experiments. (B) Temporal kinetics of LC3-II to LC3-I ratios in (A) determined using relative band densities, normalized using β-tubulin loading controls. Combined data from 4 experiments presented as mean + s.e.m for each time point compared using ANOVA with Geisser-Greenhouse correction to yield the indicated p-value. (C) Immunoblots showing LC3 expression in primary B6 mouse hepatocyte cultures inoculated with *Pb* at 24h p.i, treated as indicated from 6h. β-tubulin loading control, data representative of 2 replicate experiments. (D) LC3-II to LC3-I ratios in (C) determined using relative band densities, normalized using β-tubulin as loading controls. Data combined from separate experiments with each data point representing a replicate, compared using ANOVA with Dunnett’s correction to yield the p-values shown. (E-H) Pseudo-colored confocal images depicting the subcellular localization of LC3 in primary mouse hepatocytes infected with *Pb-GFP*, 12h p.i, (E) whole livers of B6 mice infected with *Pb-GFP,* 24h p.i. (F), primary human hepatocytes infected with *Pf*, 4d p.i (G) or whole livers of liver-humanized mice inoculated with *Pf*, 4d p.i. (H). Arrows indicate the parasite, image shown from 1 of >3 biological replicates. (I) Pseudo-colored confocal images depicting LC3 colocalization with PbUIS4 in primary B6 hepatocytes infected with *Pb*, 24h p.i. The arrow points to the parasite, representative image from 1 of 3 replicates. (J) Bar graphs depicting the frequencies of LC3 localization to the PVM (PbUIS4) in *Pb*- infected primary mouse hepatocytes 24h p.i, treated as indicated. Combined data shown from 3 replicates experiments with each point representing a replicate with >50 infected hepatocytes examined, presented as mean + s.e.m, analyzed using 1-way ANOVA withDunnett’s correction to obtain the indicated p-values. (K) Relative parasite burdens in primary B6 hepatocyte cultures infected with *Pb-Luc* and treated with IFNα/β and/or LC3siRNA, quantified using total luminescence signal at 32h p.i. Scrambled siRNA served as the control. Combined data presented as mean + s.e.m with each point obtained from a replicate experiment, analyzed using 1-way ANOVA with Tukey’s correction to obtain the indicated p-values. (L) Scatter plot indicating relative liver parasite burdens in B6 mice inoculated with *Pb* following control or LC3 siRNA delivery at −48h p.i., determined by qRT-PCR, 24h p.i. Combined data presented from three replicate experiments with each data point representing a mouse, presented as mean + s.e.m, and analyzed using unpaired t-test to obtain the indicated p-value.

Importantly, LC3 localization was evident in both *Pb* in murine hepatocytes or *Pf* in human hepatocytes, in culture or liver cryosections (**Figs 3E-H**). The strong colocalization of LC3 with PbUIS4, a marker of early PVM (Mueller et al., 2005), suggested that LC3 rapidly associates with the PV (**Figs 3I, 3J**). While the addition of exogenous IFNα/β enhanced the incidence of such association, this was reversed by NAC treatment or the absence of IFNAR (**Fig 3J**). To discount the possibility that LC3 localization to the PVM was merely delayed in the absence of IFN-I signaling, we examined co-localization of LC3 with PbHep17 that demarcates late-stage PVMs in hepatocytes (Vaughan and Kappe, 2017). We continued to observe minimal (<10%) LC3 association with *Pb* PVM even in the later stages of the infection (>32h) in the IFNARKO hepatocytes (**Figs S3B, S3C**).

Finally, to test the impact of LC3-driven LAP on IFN-I-driven *Plasmodium* control in the liver, we silenced its expression in the hepatocytes using LC3siRNA. We favored this approach instead of making conditional genetic knock-outs, considering the broad range of developmental functions carried out by LC3 (Tanida et al., 2004). Transfection of primary B6 hepatocytes with LC3siRNA resulted in significantly reduced LC3 levels but did not affect ROS levels, placing ROS upstream of LC3 following IFN-I stimulation (**Fig S3D, S3E**, **S3F**). Silencing LC3 reversed IFN-I-mediated control of *Pb* infection in isolated hepatocyte cultures (**Fig 3K**). Moreover, hydrodynamic delivery of LC3siRNA to mouse livers for reducing LC3 expression in hepatocytes - the only liver cell population productively infected by *Plasmodium* during its pre-erythrocytic development (Cowman et al., 2016) (**Fig S3G**) - greatly exacerbated parasite burdens following *Pb* Spz inoculation compared to control mice *in vivo* (**Fig 3L**). Taken together, our data indicated that LC3 activation and function induced by type-I IFNs are necessary for *Plasmodium* control in hepatocytes.

### Lysosomal control of *Plasmodium* infection downstream of IFN-I-induced ROS and LC3 in hepatocytes

A key outcome of the localization of LC3 to vacuolar membranes such as the PVM is the recruitment and fusion of lysosomes to such membranes, resulting in the acidification and destruction of the vacuolar contents (Heckmann and Green, 2019; Martinez, 2018; Wang *et al*., 2022). In line with the existing published literature (Agop-Nersesian et al., 2018; Schmuckli-Maurer et al., 2017), we observed a ‘baseline’ lysosomal association with *Pb* PVM, denoted by lysosome-associated membrane protein (LAMP)-1 colocalization with PbUIS4, in approximately 20% of the infected primary hepatocytes in culture (**Fig 4A**). The addition of exogenous IFNα/β drastically increased the frequency of lysosomal association with *Pb* PVM to >60% (**Fig 4B**), which was reversed by the NAC treatment or the absence of IFNAR (**Fig 4B**), as seen for LC3 (**Fig 3J**).

**Figure 4.**
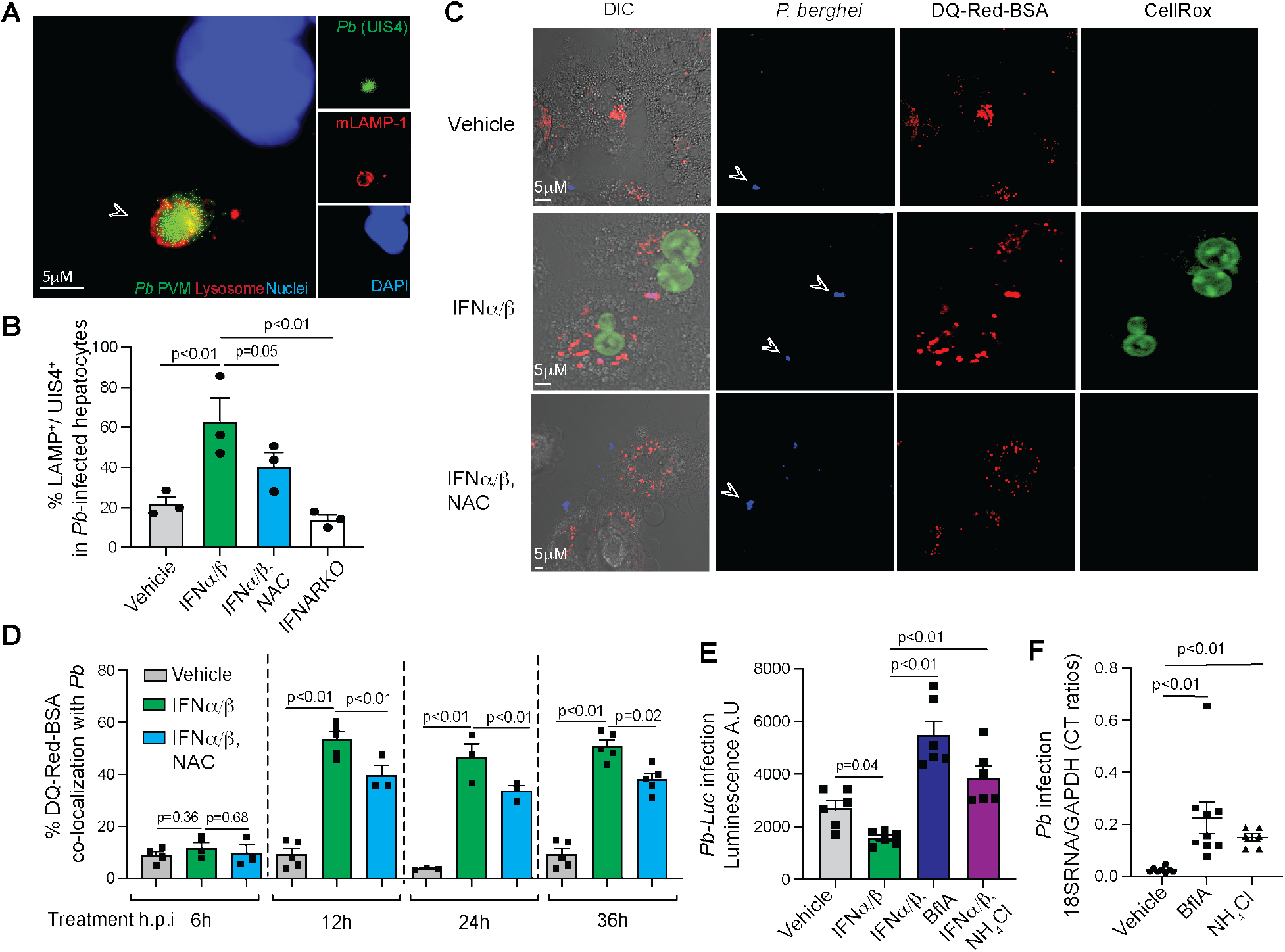
Lysosomal function instrumental in IFN-I mediated *Plasmodium* control in hepatocytes *in vitro* and *in vivo*. (A) Pseudo-colored confocal images depicting LAMP-1 colocalization with PbUIS4 in Pb- infected primary B6 hepatocytes, 24h p.i. The arrow points to the parasite, image from 1 of 3 replicate experiments with >50 infected cells/ replicate. (B) Bar graphs indicating the frequencies of LAMP-1 co-localization with PbUIS4 in primary B6 or IFNARKO hepatocytes treated as indicated at 24h p.i. Combined data from 3 separate experiments, each point represents the data from a replicate with >50 infected hepatocytes, presented as mean + s.e.m, and analyzed using 1-way ANOVA with Dunnett’s correction to obtain the indicated p-values. (C) Pseudo-colored confocal images depicting DQ-Red and CellROX signals in primary B6 mouse hepatocytes infected with *Pb* (CTV^+^), 24h p.i. following the indicated treatments from 6h p.i. Arrows indicate the parasite, image from 1 of 3 biological replicate experiments with >10 infected cells/ replicate. (D) Bar graphs indicating the frequencies of DQ-Red colocalization with *Pb* (CTV^+^) at the indicated time points from (C). Combined data shown from >3 replicate experiments with each point representing the data from a replicate with >50 infected hepatocytes analyzed, presented as mean + s.e.m and analyzed using 1-way ANOVA with Dunnett’s correction to obtain the indicated p-values. (E) Relative parasite burdens in primary B6 mouse hepatocyte cultures infected with *Pb- Luc* and treated as indicated from 6h p.i., quantified by total luminescence signal at 32h p.i. Combined data presented as mean + s.e.m from replicate experiments, analyzed using 1-way ANOVA with Tukey’s correction to obtain the indicated p-values. (F) Scatter plots indicating relative liver parasite burdens in *Pb*-inoculated B6 mice determined by qRT-PCR at 40h p.i. following treatment with bafilomycin at 0h,12h and 24h p.i. and with NH4Cl at 6h and 12h p.i. Combined data from two replicate experiments, each data point represents a mouse, presented as mean + s.e.m, and analyzed using 1-way ANOVA with Tukey’s correction to obtain the indicated p-values.

To assess the functionality of these lysosomes, we determined the level of DQ- Red and SiRlysosome staining in the *Pb* PVs. While DQ-Red is a marker of vesicular acidification, SiRlysosome indicates functional cathepsin D lysozyme (Caire et al., 2022). We observed that IFN-I not only enhanced localization but also the function of lysosomes in the *Pb*-infected hepatocytes, which was reversed by scavenging ROS or inhibiting LC3 (**Figs 4C, 4D**, **S4A-C**). Here starvation, a known inducer of lysosomal function in hepatocytes, served as a positive control (Ebner et al., 2023).

To determine whether IFN-I-enhanced lysosomal targeting and function helped control *Plasmodium*, we treated *Pb*-infected murine hepatocytes with specific inhibitors of lysosomal fusion such as bafilomycin A (BflA) or alkalinization such as ammonium chloride (NH4Cl). These treatments resulted in reduced vesicular acidification (**Fig S4D**) and reversed the IFN-I-induced control of *Pb* infection in B6 hepatocytes (**Fig 4E**). Importantly, these findings were also supported by the observation of relatively higher parasite loads in the livers of *Pb*-infected B6 mice treated with BflA or NH4Cl (**Fig 4F**). Taken together, our data suggested that lysosomes participate as direct effectors of type- I IFN-mediated killing downstream of ROS and LC3, helping accomplish *Plasmodium* control in the liver.

### IFN-I-induced GBPs combat liver-stage malaria by triggering cytosolic immunity

While our results found a critical role for IFN-I and ROS-driven lysosomal control of *Plasmodium* in the liver, various pathogen-associated molecular patterns (PAMPs) such as parasite nucleic acids are also exposed to the host hepatocyte cytoplasm, triggering additional downstream immune responses (Liehl *et al*., 2014; Marques-da-Silva *et al*., 2023). Cytosolic exposure of PAMPs such as dsDNA would invariably require the disruption of the PVM. Therefore, it was likely type-I IFNs also induce a second set of effectors that potentially disrupts the PVM for mobilizing cytosolic innate immunity.

To search for such effectors, we screened for transcriptionally enriched (>2-fold) ISGs from *Pb*-infected hepatocytes (GSE186023) that may target vesicular membranes. Several members of a single ISG family, the 65-73kDa GBPs, were highly upregulated in the infected hepatocytes *in vitro* (**Fig 5A**). Different GBPs, especially GBP1, were also transcriptionally upregulated in the livers of *Pb*-infected mice *in vivo* (**Fig 5B**). To determine which GBPs may be relevant to *Plasmodium* control in the liver, we generated tissue-specific GBPChr3KOhep and GBPChr5KOhep knockout mice, in which the GBPs present on chromosome (chr.) 3H1 (GBPs 1, 2, 3, 5, 7), or chr. 5E5 (GBPs 6, 8, 9, 10, 11) are deleted from hepatocytes using albumin-Cre excision (ms in preparation). These mice allowed us to isolate the role of GBPs in the control of *Plasmodium* in hepatocytes *in- vivo*, and how it reflected in the incidence of the resulting blood stage. GBPChr3KOhep but not GBPChr5KOhep mice showed significant deficiency in controlling *Pb* infection in the liver (**Fig 5C**). Even treatment with exogenous IFNα/β was unable to clear *Pb* infection in GBPChr3KOhep mice, causing rapid mortality, whereas *Pb*-infected GBPChr5KOhep mice exhibited partial protection following IFNα/β treatment (**Fig 5D**). These findings indicated that the GBPs encoded on Chr.3 played a crucial role in IFN-I-mediated control of *Plasmodium* in the liver *in vivo*.

**Figure 5.**
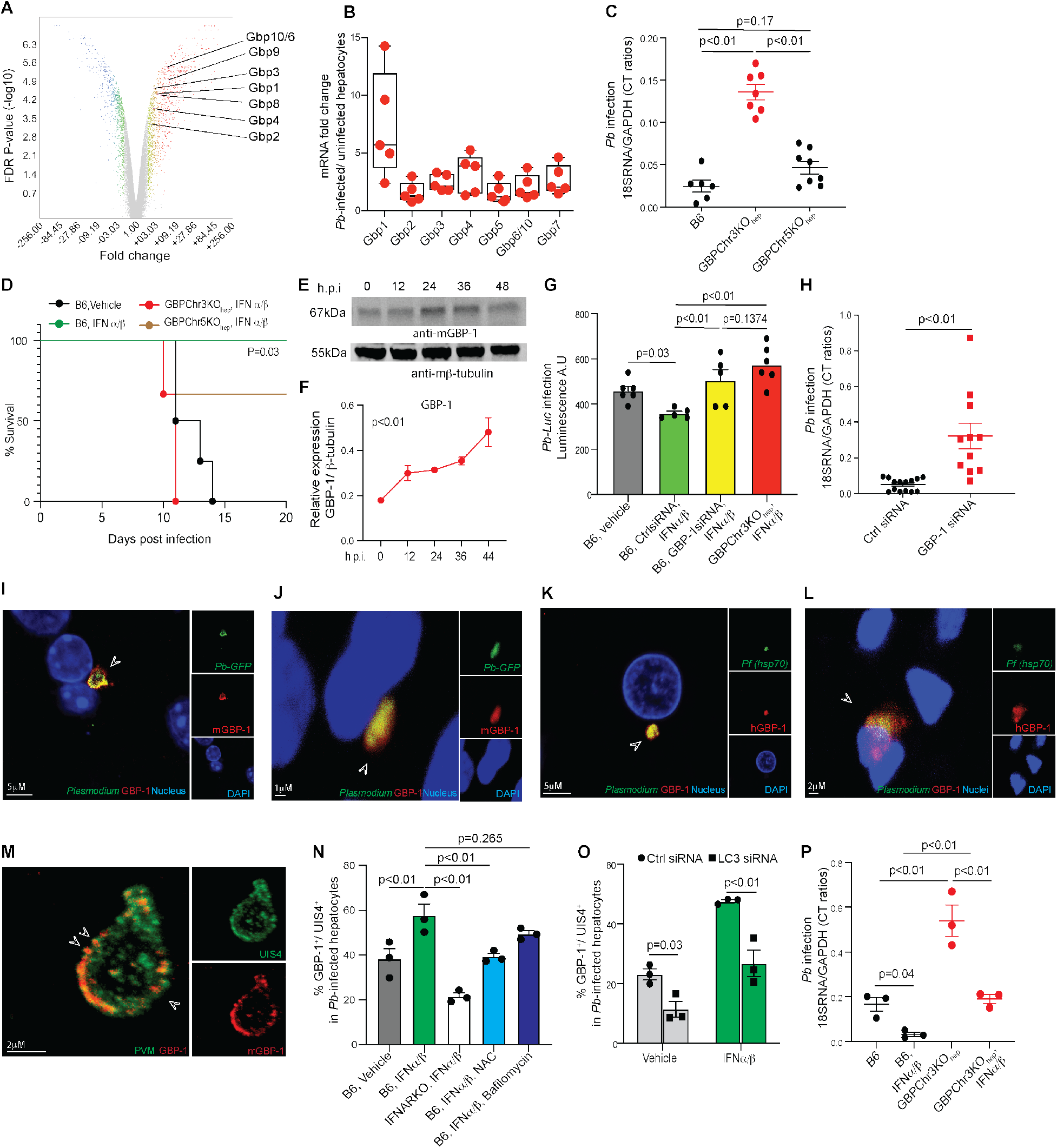
GBPs are critical for IFN I-mediated clearance of *Plasmodium* in hepatocytes *in vitro* and *in vivo*. (A) Differential expression volcano plot depicting transcriptional changes in primary B6 hepatocytes infected with *Pb* compared to uninfected hepatocytes, 36h p.i. Specific GBPs indicated. Source data: NCBI Gene Expression Omnibus GSE186023. (B) Scatter plots indicating differential expression of various GBPs in the hepatocytes isolated from *Pb*-infected (24h p.i.) versus uninfected B6 mice, determined by qRT-PCR. Representative data shown from 3 separate replicate experiments, each dot represents a separate mouse. (C) Scatter plot indicating relative liver parasite burdens in B6, GBPChr3KOhep, or GBPChr5KOhep mice inoculated with *Pb*, determined by qRT-PCR at 40h p.i. Combined data from 2 replicate experiments shown with each data point representing a mouse, presented as mean + s.e.m, and analyzed using ANOVA with Tukey’s correction to obtain the indicated p-values. (D) Survival kinetics of *Pb*-infected B6, GBPChr3KOhep, or GBPChr5KOhep mice treated with vehicle or IFNα/β at 12h and 24h p.i. Data representative of 2 separate experiments with >4 mice/group analyzed using Mantel-Cox test to yield the indicated p-value comparing GBPChr3KOhep and B6 groups. (E) Immunoblot indicating GBP1 expression in primary B6 hepatocyte cultures co- incubated with *Pb* for the indicated time-frames. β-tubulin served as loading controls. Data representative of 3 replicate experiments. (F) Temporal kinetics of GBP1 expression in the above cultures (F) quantified from relative band densities normalized to β-tubulin. Combined data presented from 4 replicate experiments presented as mean + s.e.m for each time point, compared using ANOVA with Geisser-Greenhouse correction to yield the indicated p-value. (G) Relative parasite burdens in primary mouse hepatocyte cultures infected with *Pb-Luc* and quantified by total luminescence signal at 32h p.i. following the indicated treatments. Combined data presented as mean + s.e.m, with each point representing a separate replicate, and analyzed using 1-way ANOVA with Tukey’s correction to obtain the indicated p-values. (H) Scatter plot indicating relative liver parasite burdens in B6 mice treated with control or GBP1 siRNA 24h before inoculation with *Pb*, determined by qRT-PCR at 40h p.i. Combined data from three replicate experiments, each point represents a mouse, presented as mean + s.e.m, analyzed using unpaired t-test to obtain the indicated p-value. (I-L) Pseudo-colored confocal images depicting the subcellular localization of GBP1 in primary B6 mouse hepatocytes infected with *Pb-GFP*, 24h p.i. (I), whole livers of B6 mice inoculated with *Pb-GFP*, 24h p.i. (J), primary human hepatocytes infected with *Pf*, 4d p.i (K) or whole livers of liver-humanized mice inoculated with *Pf*, 4d p.i. (L). Arrows indicate the parasite, image shown from 1 of >3 replicate experiments. (M) Pseudo-colored confocal expansion microscopy images depicting GBP1 colocalization with PbUIS4 in primary mouse (B6) hepatocytes infected with *Pb,* 24h p.i. The arrow points to what appears to be GBP1 aggregates on the PV, image representative of 10 infected cells obtained from 3 replicate experiments. (N) Bar graphs depicting the frequencies of GBP1 colocalization with PbUIS4 in *Pb*- infected *in vitro* cultured primary murine (B6 or IFNARKO) hepatocytes, 24h p.i, and treated as indicated from 6h p.i. Combined data from 3 replicate experiments, each point represents a replicate with >50 infected hepatocytes, presented as mean + s.e.m, and analyzed using 1-way ANOVA with Dunnett’s correction to obtain the indicated p-values. (O) Bar graph indicating the frequency of GBP1 colocalization with PbUIS4 in *Pb*-infected *in vitro* cultured primary murine (B6) hepatocytes at 24h p.i., following transfection with control or LC3 siRNA (-24h p.i.) and treatment with vehicle or IFNα/β at 8h p.i. Data presented as mean + s.e.m, with each point representing a replicate with >50 infected hepatocytes, analyzed using t-tests to obtain the indicated p-values. (P) Scatter plot indicating relative liver parasite burdens in B6 and GBPChr3KOhep mice inoculated with *Pb* and treated with vehicle or IFNα/β (12h, 24h p.i.) determined by qRT- PCR at 40h p.i. Combined data presented from three replicate experiments with each point representing a mouse, presented as mean + s.e.m, and analyzed using 1-way ANOVA with Tukey’s correction to obtain the indicated p-values.

Considering GBP1 was the most transcriptionally upregulated GBP in *Pb*-infected livers, and known to be a key component of cell-autonomous immunity to other intracellular protozoa such as *T. gondii* in humans and mice (Qin et al., 2017; Selleck *et al*., 2013), we examined the biology of GBP1 in the context of *Plasmodium* infection. GBP1 expression was also significantly upregulated in the *P*b-infected primary B6 hepatocyte cultures (**Fig 5E**, **5F**). We silenced GBP1 expression *in vitro* or *in vivo* using direct transfection or hydrodynamic delivery of GBP1 siRNA (**Figs S5A, S5B**). This resulted in loss of IFN-I-induced control of *Pb* infection in culture (**Fig 5G**). It also led to *Pb* infection loads similar to IFN-I-treated GBPChr3KOhep mice, suggesting that GBP1 may be the most consequential of the GBPs present on Chr.3 for *Pb* control in hepatocytes. Consist with these findings, compared to the control siRNA-treated mice, the GBP1 siRNA-treated mice exhibited significantly higher *Pb* loads in the liver (**Fig 5H**).

### GBP1 targets and disrupts the PVM within IFN-I-activated hepatocytes

GBP1 localized to *Pb* or *Pf* in both primary mouse or human hepatocytes, whether in infected cultures or liver cryosections (**Figs 5I-L**). Considering that GBP1 targets the vacuolar membrane surrounding intracellular pathogens or the outer membrane of microorganisms escaping into the cytosol with distinct immunological consequences (Kim *et al*., 2011; Praefcke, 2017; Selleck *et al*., 2013; Stickney and Buss, 2000), we examined the specific positioning of GBP1 in *Pb*-infected B6 hepatocytes, using expansion microscopy. Mouse GBP1 directly localized to the *Pb* PVM, as evidenced by its association with PbUIS4 (**Fig 5M**, **Video S1)**. We observed no association of GBP1 with the *Pb* plasma membrane, delineated by *Pb* CSP or MSP-1, in our experiments (data not shown). Consistent with these observations, GBP1 localized to the PVM of *Py*-infected hepatocytes as well (**Fig S5C**).

GBPs are known to oligomerize and mechanically disrupt their target membranes through conformational extension (Weismehl *et al*., 2024; Zhu *et al*., 2024). To investigate the potential role of GBP1 in disrupting the PVM, we selectively permeabilized the plasma membrane, but not the PVM in primary *Pb*-infected hepatocytes cultures derived from GBPChr3KOhep or AIM2KO mice, using 0.01% digitonin (Niklas et al., 2011). We then employed CSP-1 antibodies that mark the plasma membrane of *Pb* schizonts to ascertain the presence of an intact PVM. In this assay, undamaged PVMs exclude antibodies from accessing the parasite’s plasma membrane. AIM2KO mice were used for comparison to prevent pyroptotic elimination of the host cells following the potential loss of PVM integrity, and CSP was chosen as the target to detect the parasite in the early stage of the infection; IFN-I treatment would clear the infection rapidly with time. Indeed, the absence of GBPs significantly impeded the CSP-1 antibodies from accessing *Pb* in the majority of the infected hepatocytes (denoted by the presence of Hep17 or CSP), indicating that GBPs are needed for disrupting the PVM (**Figs S5D, S5E**).

These findings were further supported by high-resolution confocal imaging showing disintegrating *Pb* parasites that were associated with GBP1, within the B6 hepatocytes (**Fig S5F**). An internal structural protein of the parasite, acyl carrier protein (ACP) (Gallagher and Prigge, 2009), appeared to be discharged following GBP1- mediated disruption of the PV that possibly occurs in parallel with lysosome-mediated destruction of the parasite. Lysosome-mediated destruction of *Plasmodium* coupled with GBP-mediated disruption of its PVM result in the exposure of *Plasmodium* antigens and PAMPs to the hepatocyte cytoplasm.

We tested if GBPs 2 or 5, which are present on Chr.3 and reported to control intracellular pathogens like *Chlamydia*, *Francisella*, *Shigella,* and *T. gondii* (Kravets et al., 2016; Lindenberg et al., 2017; Matta et al., 2018; Meunier et al., 2015) also contributed to *Plasmodium* control in the liver. Both GBPs 2 and 5 failed to localize to *Pb* in the infected hepatocytes (**Figs S6A, S6B**) and silencing their expression in hepatocytes using hydrodynamic delivery of siRNAs (**Fig S6C, S6D**) did not impact *Pb* burdens in the livers of mice (**Fig S6E**). This differs from the closely related apicomplexan parasite, *T. gondii*, where murine GBP2 and GBP5 can target the parasitophorous vacuole (Degrandi et al., 2013; Kravets *et al*., 2016).

Notably, *Pb*-infected hepatocytes in culture exhibiting GBP1 localization to the PVM was enhanced by IFN-I treatment, and dependent on IFNAR, ROS, and LC3, but independent of lysosomal function (**Figs 5N, 5O**). IFN-I treatment still significantly diminished the *Pb* infection loads in GBPChr3KOhep mice, although it was not to the extent observed in B6 mice (**Fig 5P**). This implied that while GBPs are important, they are not the sole effectors of the IFN-driven control of liver-stage malaria and that lysosomal killing is also required.

### GBP1 is recruited downstream of ROS and LC3 targeting of *Plasmodium* PVM

GBP deficiency did not impact IFN-I-induced ROS production nor lysosomal association with *Pb* (**Fig S7A, S7B**), whereas ROS was necessary for GBP1 localization to the PV (**Fig 5N**). Thus, GBP1 probably operates downstream of ROS production and parallel with lysosomal killing. Indeed, GBP1 localization to *Pb* PVM occurred independently of lysosomal acidification and proteolysis (**Fig 5N**), indicating that GBP and lysosomal function constitute two critical yet independent consequences of IFN-I signaling in the *Plasmodium*-infected hepatocytes. While inhibiting lysosomes was insufficient to fully reverse IFN-I-induced inhibition of *Pb* in B6 mice, it did so effectively in the GBPChr3KOhep mice (**Figs S7C, S7D**). This finding supports a dual defense strategy in which lysosomes and GBPs act as non-redundant and complementary effectors to eliminate *Plasmodium*, with the former likely lysing and the latter damaging the PV to promote a parasiticidal environment.

Notably, LC3 silencing interfered with GBP1 targeting to the PVM (**Fig 5P**), as reported for *T. gondii*-infected cells (Haldar et al., 2015; Park et al., 2016) . We generated a transgenic mouse hepatocyte line (Hepa1-6) which polycistronically expressed LC3 and GBP1, tagged respectively with red or green fluorescent proteins (LC3-RFP, GBP1-GFP). LC3 initially co-localized with *Pb* before more persistent association of GBP1 (**Video S2, Figs S8A, S8B)**. While LC3 was required for GBP1 association with the PVM (**Fig 5O**), LC3 still converged on *Pb* in GBPChrKOhep hepatocytes (**Fig S8C)** at frequencies similar to B6 hepatocytes (data not shown). Furthermore, while LC3 knockdown resulted in higher liver parasite loads in *Pb*-infected mice, the additional knockdown of GBP1 did not further enhance parasite burdens (**Fig S8D**). These data indicated that LC3 activation and function along with ROS occurred upstream of GBP1 localization to the PV in *Plasmodium*-infected hepatocytes and is a prerequisite for GBP1 function against malaria.

### GBP1 lysis of the PVM facilitates caspase-1 processing in *Plasmodium*-infected hepatocytes

GBP1-mediated lysis of PVMs in IFN-I-activated hepatocytes may release ligands for activation of cytosolic innate immune pathways. Indeed, we reported earlier that the cytosolic receptor, AIM2, detects *Plasmodium* dsDNA in the cytosol and drives caspase- 1 inflammasome-mediated gasdermin D (GSDMD) activation, resulting in pyroptosis of hepatocytes (Marques-da-Silva *et al*., 2023; Marques-da-Silva *et al*., 2024a). A frequent feature of AIM2 inflammasomes is their close proximity to intracellular pathogens, including *Plasmodium* in hepatocytes (Fernandes-Alnemri et al., 2010; Marques-da-Silva *et al*., 2023).

Considering that GBPs can help assemble caspase-1 complexes and interact with other caspases to regulate their function in a variety of models (Fisch *et al*., 2020; Shenoy *et al*., 2012; Wandel *et al*., 2020; Zhu *et al*., 2024), we hypothesized that GBP1 may directly facilitate caspase-1 inflammasome formation in *Plasmodium*-infected hepatocytes as well. Caspase-1 and GBP1 exhibited marked colocalization with *Pb* or *Pf* in the vast majority of primary mouse and human hepatocytes (**Figs 6A-D**). Confocal imaging showed distinct convergence of GBP1 and caspase-1 at the PVM (**Fig 6E**). To test if endogenous GBP1 and caspase-1 interacted under these conditions, we immunoprecipitated caspase-1 from the *Pb*-infected primary hepatocyte cultures and probed elutes using a GBP1-specific antibody. Endogenous GBP1 co- immunoprecipitated with caspase-1 (**Fig 6F**). These findings revealed that GBP1 converges with and binds caspase-1-containing complexes in *Plasmodium*-infected hepatocytes.

**Figure 6.**
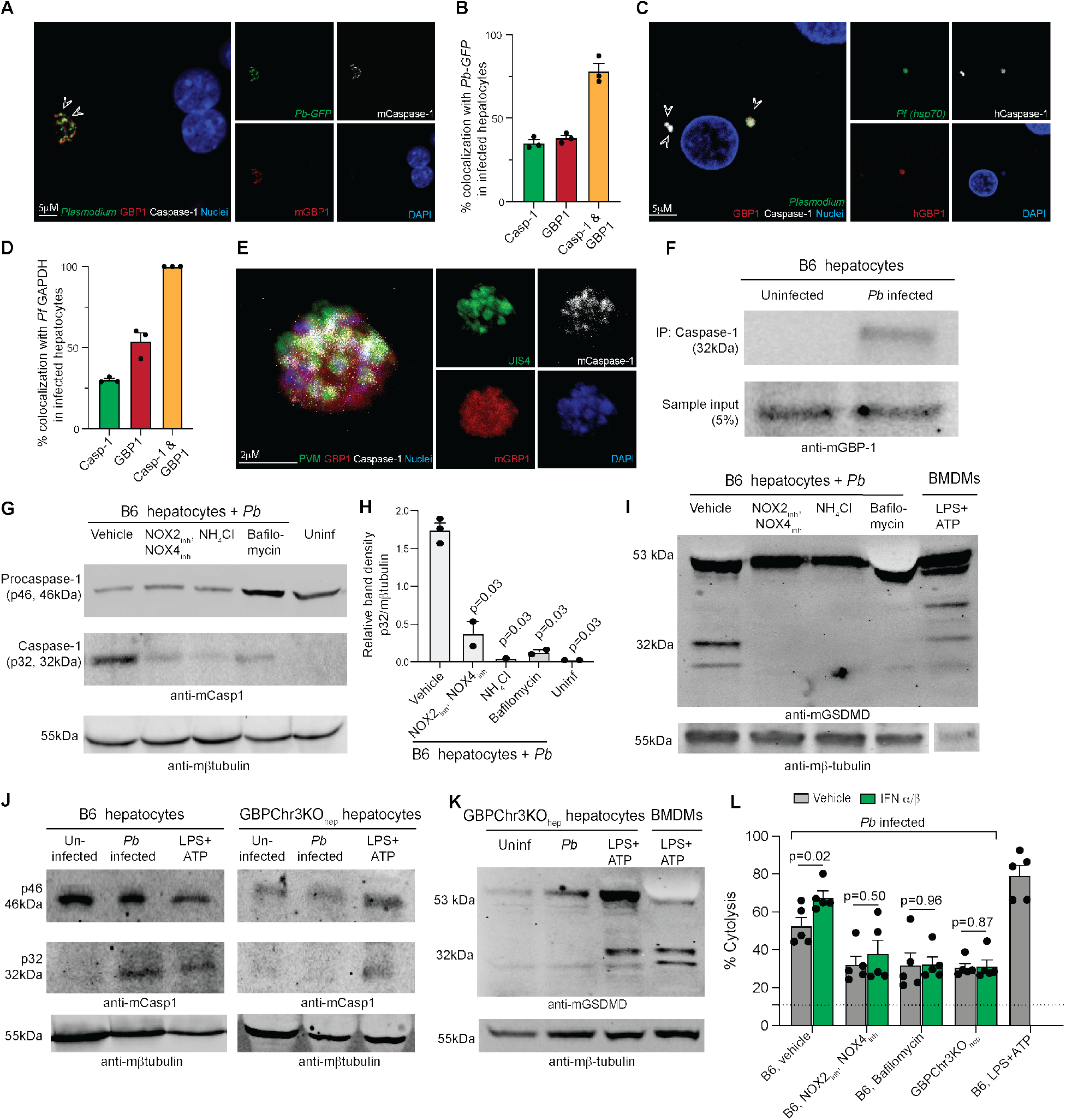
Endogenous GBP1 associates with caspase-1 and promotes pyroptosis in *Plasmodium*-infected hepatocytes. (A) Pseudo-colored confocal images depicting GBP1 and caspase-1 localization in *PbGF*P-infected primary murine (B6) hepatocytes, 36h p.i. The arrows points to the foci of GBP1 and caspase-1 colocalization. Representative image from 1 of 3 replicate experiments with >10 infected cells analyzed/sample. (B) Bar graph indicating the frequency of GBP1 and caspase-1 colocalization with *Pb- GFP* in the infected primary B6 hepatocytes, 36h p.i. Combined data shown from 3 replicate experiments, each point represents a replicate with >50 infected hepatocytes, presented as mean + s.e.m. (C) Pseudo-colored confocal images depicting GBP1 and caspase-1 colocalization with Pfhsp70 in *Pf*-infected primary human hepatocytes, 4d p.i. The arrows point to *Pf.* Representative image from 1 of 3 replicate experiments with >10 infected cells analyzed/sample. (D) Bar graph indicating the frequency of GBP1 and caspase-1 colocalization with *Pf* in primary human hepatocytes, 4d p.i. Combined data shown from 3 replicate experiments, each point represents a replicate with >50 infected hepatocytes evaluated, presented as mean + s.e.m. (E) High-resolution confocal images depicting GBP1 and caspase-1 co-localization with PbUIS4 in *Pb*-infected primary murine (B6) hepatocytes, 36h p.i. Representative image from 1 of 3 replicate experiments with >10 infected cells analyzed/replicate. (F) Immunoblot analysis for GBP1 co-immunoprecipitated with caspase-1 in whole-cell lysates of primary mouse (B6) hepatocyte cultures infected or not with *Pb*, at 30h p.i. Sample input served as loading control. Data represent 2 separate replicate experiments. (G) Immunoblot indicating p46 and p32 caspase-1 in primary B6 hepatocyte cultures infected with *Pb*, 30h p.i., following treatments with the indicated reagents. Vehicle treated or uninfected hepatocytes served as controls; β-tubulin served as loading controls. Data represents 4 separate replicate experiments. (H) Caspase-1 p32 in (G) quantified using band densities normalized to β-tubulin. Each data point represents a replicate experiment, analyzed using ANOVA with Dunnett’s correction comparing each group to the vehicle control group yielding the indicated p- value. (I) Immunoblot depicting GSDMD processing in primary B6 hepatocyte cultures infected with *Pb*, 30h p.i. following pre-treatments with the indicated reagents. BMDMs treated with LPS+ATP served as treatment control and β-tubulin served as loading control. Data represents 2 separate replicate experiments. (J) Immunoblot indicating p46 and p32 caspase-1 in primary B6 or GBPChr3KOhep hepatocyte cultures infected with *Pb*, 30h p.i., or treated with LPS+ATP (16h). Uninfected hepatocytes served as additional controls; β-tubulin served as loading controls. Data represents 3 separate replicate experiments. (K) Immunoblot depicting GSDMD processing in uninfected, *Pb*-infected (30h p.i.) or LPS+ATP treated primary GBPChr3KOhep hepatocyte cultures. GBPChr3KOhep BMDMs treated with LPS+ATP served as treatment control and β-tubulin served as loading control. Data represent 3 separate replicate experiments. (L) Cytolysis determined by LDH release in cultured primary mouse hepatocytes from B6 or GBPChr3KOhep mice, infected with *Pb* at 36h p.i. following treatment with the indicated reagents from 6h p.i. The dotted line indicates uninfected B6 hepatocyte culture. Combined data presented with each point representing a replicate experiment, presented as mean + s.e.m and analyzed using unpaired t-tests to yield the indicated p-values.

### GBP1 drives IFN-I-induced pyroptosis in *Plasmodium*-infected hepatocytes

Based on the above results, lysis of *Plasmodium* and its PV by type-I IFN-driven responses including GBPs may trigger caspase-1 activation within infected hepatocytes. Blocking NOXs or lysosomal function significantly impeded caspase-1 activation and GSDMD processing in the *Pb*-infected B6 hepatocytes (**Figs 6G-I**). Similarly, caspase-1 and GSDMD activation was undetectable in *Pb*-infected GBPChr3KOhep hepatocytes (**Fig 6J,K**). The absence of caspase-1 and GSDMD activation coincided with reduced cell death compared to the vehicle-treated *Pb*-infected or LPS+ATP-treated control B6 cells (**Fig 6L**). Considering that the absence of lysosomal function alone in the GBP-sufficient B6 hepatocytes partly impeded caspase-1/GSDMD activation suggests that lysosomes and GBPs operate together - the former by lysing *Plasmodium* and the latter by damaging the PV - to induce pyroptosis in *Plasmodium*-infected hepatocytes (**Fig S9**). Hence these two hepatocyte pathways co-operate to mobilize vacuolar and cytosolic immunity against malarial infection *in vitro* and *in vivo*.

## Discussion

Type-I IFN mobilizes cell-autonomous defense in non-immune cells such as hepatocytes infected with *Plasmodium*; however, the inhibitory mechanisms remain unknown (Gazzinelli *et al*., 2014; He *et al*., 2020; Marques-da-Silva *et al*., 2024b). In this study, we found type-I IFNs induce NOX2- and NOX4-mediated ROS production, LAP, as well as GBP1 expression and localization to the *Plasmodium* PV in infected human and murine hepatocytes. LC3 decoration of PVM led to lysosomal and GBP1 fusion with it, resulting in the disruption of the PV. GBP1 also appeared to anchor the caspase-1 inflammasome to promote hepatocyte pyroptosis (**Fig S9**). Importantly, these mechanisms operated within an intact mammalian host as well.

*Plasmodium* dsDNA and RNA have been shown to gain access to the hepatocyte cytoplasm during the liver stage of malaria and trigger downstream immune responses (Liehl *et al*., 2014; Marques-da-Silva *et al*., 2023; Marques-da-Silva *et al*., 2024a). How nucleic acids of *Plasmodium,* concealed behind the PVM and plasma membrane, would gain access to the host cell cytoplasm has remained an outstanding question in the field. IFN-I-induced lysosomal destruction of *Plasmodium* coupled with the GBP-mediated disruption of the PVM appears to offer some explanation.

Autoproteolytic caspase-1 processing is dependent on achieving a threshold concentration in the cytoplasm (Boucher et al., 2018; Walsh et al., 2011). Assembly of procaspase-1 complexes may enable it to surmount this threshold and undergo cleavage in hepatocytes, considering its inherently reduced expression in those cells (Marques-da- Silva *et al*., 2024a). GBP1 appears critical in supporting the inflammasome pathway in hepatocytes by enabling pro/caspase-1 complex formation at the PVM, like it does in *Salmonella*-infected epithelial cells with caspase-4 (Wandel *et al*., 2020; Zhu *et al*., 2024). This would expose *Plasmodium* PAMPs to the hepatocyte cytoplasm to instigate innate immune signaling and further support it by helping assemble the inflammasome complex.

A key unresolved question is why both lysosomal and GBP functions are enlisted independently to eliminate *Plasmodium*. What might be the purpose of such duality? Lysosomal killing may directly eliminate the parasite, whereas GBP1-mediated disruption of the PVM also exposes *Plasmodium* PAMPs for cytosolic innate sensing and *Plasmodium* antigens for cytosolic MHC class I presentation by the liver cells. Indeed, blocking caspases prevents antigen-presenting cells (APCs) from efficiently acquiring *Plasmodiu*m antigens from the liver following *Py* or *Pb* Spz infections (Kurup *et al*., 2019a; Marques-da-Silva *et al*., 2024a). These are necessary to generate protective CD8 T cell responses during natural infections and to Spz-based malaria vaccines (Kurup et al., 2019b). Our work thus offers a bridge between IFN-I mediated cell-intrinsic immunity and CD8 T cell-mediated adaptive responses against liver-stage malaria.

Existing treatments against malaria primarily target the parasites in their blood stage, where the parasite numbers are greatest. This creates selection pressure that likely hastens the emergence of new drug-resistant parasites. The constant emergence of new drug-resistant lines is one of the major challenges we are facing in combating malaria today (Conrad and Rosenthal, 2019). With significantly fewer parasite numbers, the liver stage offers an attractive opportunity for anti-malarial drug and immune vaccine development (Antonova-Koch et al., 2018). Host innate immune pathways can be potentially manipulated to limit *Plasmodium* infection to the liver itself and minimize emergence of new drug resistance. Such drugs would be attractive candidates for prophylactic administration in malaria-endemic regions (von Seidlein and Greenwood, 2003). Immunotherapeutic drugs or adjuvants that augment innate immune responses in the liver could also help clear dormant *P. vivax* infections (hypnozoites) that seed relapsing blood-stage malaria. IFN-α treatment was recently shown to clear hypnozoites in human hepatocytes (Mancio-Silva et al., 2021), indicating that immunotherapeutic approaches could theoretically help prevent malaria relapses. Such approaches have precedent with recombinant type-II IFN being clinically used to clear numerous human pathogens (Casanova et al., 2024), as first shown for lepromatous leprosy (Nathan et al., 1986). The application of IFN-I-induced innate immunity, however, has not been explored to the same extent but our knowledge of host defense against parasitic infections has continued to grow over the past decade (Hunter and Sher, 2014).

Mobilizing non-immune cells via IFN signaling to clear malarial parasites reinforces the idea that cells outside of the hematopoietic compartment exert underappreciated roles in host defense (Gaudet et al., 2021). This idea is coupled with the fact that IFNs often represent a potent signal for protective immunity during malaria infection in humans. People with higher serum IFN levels consistently exhibit significantly reduced incidence and recurrence of malaria in endemic areas (Deloron et al., 1991; Krupka et al., 2012; Luty et al., 1999; Subramaniam et al., 2015). Elevated serum IFN-α levels in children coincide with milder disease, and disruptive polymorphisms in *IFNA* promoters associate with increased malarial susceptibility and mortality (Kempaiah et al., 2012; Luty et al., 2000). Our work in identifying some of the critical effectors against *Plasmodium* infection in primary hepatocytes offers a solid framework for future studies attempting to therapeutically modify IFN-I-driven immune responses in the liver.

## Acknowledgements

We would like to thank Drs. Fernando Sanchez-Valdez (UGA), Drew Etheridge (UGA), Rahul Vijay (RFU), and Photini Sinnis (Johns Hopkins) for technical assistance, the UGA CTEGD Flow Cytometry Core, UGA CVM Cytometry Core facility, UGA CTEGD Sporocore, and UGA CTEGD animal research facility staff for their support. These studies were funded by the US National Institutes of Health (T32AI060546 to CSS and JS, AI153290 to DEK, AI114543, AI167847, AI178159, AI185067, AI185725 to JTH, R01AI068041-15, R01AI108834-10 to JDM and R01AI168307 to SPK). DEK is a Georgia Research Alliance scholar, JDM is an Investigator of the Howard Hughes Medical Institute.

## Author contributions

Conceptualization: CMdS, SPK; methodology: CMdS, CSS, CB, JCS, SPK; investigation: CMdS, CSS, CB, ECC, JCS, DEK, ESP, ZY, PHK, JTH, JDM, SPK; writing: CMdS, JTH, JDM, SPK; funding acquisition: DEK, JTH, JDM, SPK; supervision: SPK.

## Video S1: GBP1 colocalizes with PbUIS4

High-resolution expansion immunofluorescence microscopy and 3D volumetric reconstruction of *Pb*-infected B6 hepatocyte showing the relative distribution of PbUIS4 (pseudocolored green) and GBP1 (pseudocolored red), 24h p.i. DNA stained with DAPI (pseudocolored purple). Video representative of 10 infected cells obtained from 3 separate experiments. Green: PbUIS4, Red: GBP1, Purple: DAPI

## Video S2: LC3 precedes GBP1 localization to *Pb*

Confocal time-lapse live-microscopy images depicting the kinetics of localization of LC3- RFP and GBP1-GFP with *Pb* (CTV^+^) in transgenic Hepa1-6LC3-GBP1 reporter hepatocytes. Images obtained 10 minutes apart for a total of 60 minutes. Video representative of >5 infected cells from 2 separate experiments.

## Methods

### Mice and murine pathogens

C57BL/6 (B6) and IFNAR1KO mice were purchased from the Jackson Laboratory. Liver-humanized mice were procured from Yecuris. We generated the GBPCr3KOhep and GBPCr5KOhep mice. All mice were housed with appropriate biosafety containment at the University of Georgia or Johns Hopkins University animal care units. The animals were treated and handled in accordance with the guidelines established by the respective Institutional Animal Care and Use Committees. *A. stephensi* mosquitos infected with *P. berghei ANKA* (*Pb*), *Pb* expressing luciferase (*Pb-Luc*) or *Pb* expressing GFP (*Pb-GFP*) were reared at the Sporocore at the University of Georgia.

### Tertiary cell culture and transgenesis

Mouse Hepa1-6 hepatoma cells (ATCC) were cultured in Dulbecco’s Modified Eagle Medium (DMEM, Gibco/Invitrogen) supplemented with 10% fetal calf serum (FCS) (Sigma-Aldrich), 100 U/ml penicillin and 100 µg/ml streptomycin (Gibco/Invitrogen). Cells were maintained in a humidified cabinet at 37°C and 10% CO2. To generate transgenic Hepa1-6 cells, 1×10^6^ cells were pelleted by centrifugation. After resuspension in Nucleofector V solution (Lonza, VVCA-1003), the cells were electroporated with 3µg of the GBP1-eGFP-LC3-RFP DNA (Vector Builder) by using the program T-28 on the Nucleofector 2b transfection device according to the manufacturer’s instructions. The cells were then seeded onto glass bottom dishes for live cell imaging.

### Murine primary hepatocyte culture, *in vitro* sporozoite infection, and treatments

Primary hepatocytes were isolated from mice as described in detail before (Marques-da- Silva *et al*., 2023). In short, in anesthetized (Ketamine/Xylazine, 87.5/12.5 mg/Kg) (Sigma-Aldrich) mice, the inferior vena cava was catheterized (BD auto guard, 22G) aseptically to perfuse the liver by draining through the portal vein. Steady-state perfusion of the liver was performed first with PBS (4ml/min for 5 minutes), then Liver Perfusion Medium (4ml/min for 3 minutes, Gibco), and finally the Liver Digest Medium (4ml/minute for 5 minutes, Gibco). The digested liver was excised, single cell suspension made and resuspended in DMEM (Gibco/Invitrogen) with 10% FCS (Sigma-Aldrich). Hepatocyte fraction was recovered by centrifugation at 57g, from which debris and dead cells were removed by density gradient centrifugation with 46% Percoll (GE Healthcare). The pelleted cells were counted and resuspended in DMEM with 10% FCS. Cells were plated on collagen-coated plates or dishes and incubated at 37°C, 5% CO2. Primary hepatocyte cultures were inoculated with unless otherwise specified, 1 Spz per 4 hepatocytes. Cultures were further incubated for the desired time to allow infection and liver-stage parasite development. Unless otherwise specified, hepatocyte cultures were treated from 6h with 20mM N-acetyl cysteine (Sigma-Aldrich), 50nM Bafilomycin (ChemCruz), 20mM NH4Cl (Sigma-Aldrich), 1mM GSK2795039 (Tocris) or 10µM GLX35122, (MedChem Express) or murine type-I IFNs (100U/mL of IFNα and 100U/mL IFNβ, PBL). Culture supernatants and/or cells and their lysates were obtained at various time points post-infection or stimulation as indicated. Starvation controls followed previous protocols: cells were rinsed 3 times with PBS and subsequently incubated with EBSS (0.001M calcium chloride, 0.00081M magnesium sulfate heptahydrate, 0.005M potassium chloride, 0.026M sodium bicarbonate, 0.117M sodium chloride, 0.001M sodium phosphate monobasic monohydrate and 0.006M D-glucose) at 37 °C, 5% CO2 for a minimum of 6h.

### *Plasmodium* inoculations into mice

For sporozoite challenge experiments, salivary glands of *Pb-*parasitized *A. stephensi* mosquitoes were dissected, and the sporozoites were isolated, counted, and injected 2x10^4^ in 200μL RPMI (Sigma-Aldrich) with 1% mouse serum (Innov-research) into the tail vein of mice as described before (Kurup *et al*., 2019a). For infecting liver-humanized mice, *A. stephensi*-infected with *P. falciparum* NF54 strain were allowed to bite ketamine-anesthetized liver-humanized mice for 15 mins. The remaining sporozoites from the same mosquitoes were isolated and inoculated intravenously into the tail vein of the mosquito- fed mouse at 7.5x10^5^ sporozoites per mouse.

### Primary human hepatocyte culture, *Pf* infection, and treatments

Primary human hepatocyte infection in this study followed published methods (Marques-da-Silva *et al*., 2023). Briefly, cryopreserved primary human hepatocytes (BioIVT) were thawed two-days before infection and seeded with 18x10^3^ cells per well in a 384-well collagen-coated plate (Greiner). Hepatocytes were maintained with daily media change using customized InVitroGro HI medium devoid of dexamethasone (BioIVT) supplemented with 5% human serum (Interstate Blood Bank), penicillin, streptomycin, neomycin, and gentamicin (Gibco). The salivary glands of *Pf* BD007 (Bangladesh isolate)- infected *A. stephensi* mosquitoes were collected, total sporozoite counts were determined and 6x10^3^ sporozoites were inoculated into the hepatocyte culture. The hepatocytes in culture were maintained at 38°C with 5% CO2. *Pf*-infected hepatocytes were treated or not with 10 mM N-acetyl cysteine, 100 µM GSK2795039 (NOX2 inhibitor, Tocris), 10µM GLX35122 (NOX4 inhibitor, MedChem Express) or stimulated human type 1 IFNs (100U/mL IFNα and IFNβ, PBL) as indicated. Medium with/out the treatments was changed daily until 4 dpi.

Immunofluorescence assays were performed at 4d p.i. Infected hepatocytes were fixed with 4% paraformaldehyde (Alfa Aesar) for 10 min at room temperature and washed twice with PBS followed by permeabilization and blocking with 0.03% Triton-X (Acros) and 1% BSA (w/v) (Fisher Scientific) in PBS. The samples were concurrently stained with anti-GAPDH and anti-MSP antibodies (European Malaria Reagent Repository, UK) at 4°C overnight followed by staining with fluorescently labeled secondary antibodies at 4°C, overnight. The wells were then stained with DAPI (Sigma-Aldrich) at room temperature for 1h. The *Pf* infection in each well was imaged using ImageXpress and quantified using the MetaXpress software (Molecular Devices).

### Assessment of parasite burden

Liver parasite burden was assessed by quantitative real-time PCR for parasite 18S rRNA in hepatocytes derived from mice, as described in detail before (Doll et al., 2016). In short, total RNA was extracted from the liver at the indicated time points after *Plasmodium* inoculation using TRIzol,(Sigma-Aldrich) followed by DNase digestion/cleanup with RNA Clean and Concentrator kit (Zymo Research). 2μg liver RNA per sample was used for qRT-PCR analysis for *Plasmodium* 18S rRNA using TaqMan Fast Virus 1-Step Master Mix (Applied Biosystems). Data were normalized for input using GAPDH control in hepatocytes for each sample and are presented as ratios of *Plasmodium* 18S rRNA to GAPDH RNA. The ratios depict relative parasite loads within an experiment and do not represent absolute values.

Alternatively, *in vivo*, *Pb-Luc* was used to assess the kinetics of replication and clearance of *Plasmodium* infection in the liver of mice. For bioluminescence detection, mice were injected with D-luciferin (150 mg/kg; PerkinElmer) intraperitoneally and anesthetized using 2% (vol/vol) gaseous isoflurane (Dechra) in oxygen before imaging on an IVIS 100 imager (Xenogen). Quantification of bioluminescence and data analysis was performed using Living Image v4.3 software (Xenogen).

*Pb-Luc* infection loads were determined *in vitro* using Bright-Glo (Promega) as described before (Derbyshire et al., 2012). In brief, primary hepatocytes were plated for 24h on a 96-well collagen-coated plate. Then, the cells were infected with *Pb-Luc,* and 6h post-infection (hpi), the cells were treated or not with the specified inhibitors. 8 hpi, cells were treated with murine IFN-α and IFN-β for an additional 24 hours. Finally, Bright-Glo reagent was added to evaluate parasite loads (as indicated by manufacturer’s). The relative signal intensity of each plate was evaluated with Spectramax iD3 (Molecular Devices) and the SoftMax Pro 7.1.2 software.

### Assessment of gene expression

Host gene expression was determined in hepatocytes isolated from whole livers. Total RNA was extracted from the cultured or freshly isolated hepatocytes using the TRIzol reagent (Sigma-Aldrich), followed by DNase digestion/cleanup with RNA Clean and Concentrator kit (Zymo Research). 2μg liver RNA per sample was used for qRT-PCR analysis for various genes using specific primers (GBPs (Meunier *et al*., 2015), NOX2, or NOX4 (Sancho et al., 2012; Wan et al., 2022). Data were normalized against GAPDH gene transcripts in hepatocytes (Marques-da-Silva *et al*., 2022).

### Transcriptome analysis

Hepatocytes, infected with *Pb-GFP* or naïve (from uninfected cultures) prepared as previously described were FACS sorted and re-suspended in TRIzol. RNA was then purified using the RNAeasy kit (Qiagen). RNA purity and quality were assessed with an Agilent 2100 Bioanalyzer. Transcriptome analysis was conducted on the Affymetrix GeneChip Mouse Transcriptome array 1.0. The resulting data were analyzed and visualized using Affymetrix Expression Console v1.4.4.46, Applied Biosystems Transcription Analysis Console v4.0.1.36, and Interferome v2.01 (www.interferome.org), and subsequently submitted to the NCBI Gene Expression Omnibus (GSE186023).

### Microscopy

Liver sections collected from infected (5x10^5^ *Py* or 7.25x10^5^ *Pf* sporozoites) mice were fixed, permeabilized with 1% Triton X-100 (Fisher Bioscience) and imaged after staining. Images were acquired on SP8 NLO Microscope (Leica) using a 10X/0.40 dry objective (live) or 25x/0.95 water immersion objective with coverslip correction (fixed), as described in detail previously (Kurup *et al*., 2019a; Kurup et al., 2017). All images acquired were processed using Imaris software (Bitplane).

Pfhsp70 (polyclonal, GenWay) was used to mark *Pf.* The following were used to identify *Pb*: UIS4 (gift from Dr. Scott Lindner, Pennsylvania State University), ACP (gift from Dr. Stefan Kappe, Seattle Children’s Hospital), or CSP (this paper). For live imaging, hepatocytes were cultured overnight in collagen-coated chamber slides (ibidi) and infected with *Plasmodium*. Cultures were maintained in a climate-controlled chamber during imaging.

Primary murine and human hepatocytes were attached to collagen (Sigma- Aldrich)-coated slides for 24h. Slides were then infected for 24-36h with *Pb,* or for 4 days with *P. falciparum*. Slides were washed twice with PBS and fixed with 4% paraformaldehyde (Alfa Aesar)/PBS (20 min), permeabilized with Cytofix/Cytoperm buffer (BD Biosciences) for 15 min/ 4°C or 1% tritonX100 (Acros), washed once with Perm/Wash buffer (BD Biosciences), treated with Cytoperm Permeabilization Buffer Plus (BD Biosciences), for 15 min/ 4°C, washed again with Perm/Wash buffer, treated again with Cytofix/Cytoperm (BD Biosciences) for 15 min/ 4°C. Samples were co-incubated with rabbit monoclonal anti-UIS4 antibody (In house generated), anti- mouse LC3 (Clone 4E12, MBL International), anti- mouse Lamp 1 (Clone 5E7, ABclonal), anti-mouse GBP1 (Clone 1B1, Santa Cruz), anti-mouse galectin 3 (polyclonal, R&D systems), anti-mouse Rubicon (polyclonal, Abcam), anti-mouse NOX2 (Clone 53, BD Biosciences), anti-mouse NOX4 (Clone C3, Santa Cruz), anti-CSP (sporozoites, this paper), anti-ACP (schizont, Stephan Kappe) or anti-*Pf* GAPDH (European Malaria Reagent Repository, UK) antibodies. Subsequently, the samples were washed thrice with Perm/Wash buffer, probed with 5 mg/mL fluorophore-conjugated secondary antibodies and 0.1 mg/mL of DAPI (Sigma-Aldrich), washed thrice with Perm/Wash buffer, mounted in Vectashield mounting medium (Vector) and imaged. Images were acquired on SP8 NLO (Leica) or Zeiss LSM880 and processed using Imaris software (Bitplane).

Co-localization was quantified by analyzing random fields, identifying at least 50 infected cells (UIS4^+^ or GFP^+^) from at least 3 separate cultures. Quantification was performed visually using the microscope. If the signal covered >30% of the area/ perimeter of the parasite, it was considered positive.

Lysosomal staining was performed in live primary hepatocytes plated in 35mm^2^ collagen-coated dishes. *Pb* stained with cell trace violet (*Pb*-CTV, Thermofisher) was inoculated for 6h, treated or not with NAC (Sigma-Aldrich) for an additional 2h, then treated with vehicle (PBS) or type-I IFNs (PBL) for the indicated time-points. Dishes were incubated with DQ-Red BSA (InVitrogen) or SiR-Lysosome (Spirochrome) according to manufacturer’s protocol. In brief, at the indicated time-points, cells were washed twice with PBS, then incubated for 1h at 37°C with DQ-Red BSA (10 µg/mL in DMEM + 10% FBS) or SiR-Lysosome (1 µM in DMEM + 10% FBS), washed twice with PBS and imaged. Images were acquired Zeiss LSM880 with Zen 2.3 imaging software and images processed using Imaris software (Bitplane).

Expansion microscopy was performed as described before (Atchou et al., 2023; Liffner and Absalon, 2021). In short, coverslips with 4% PFA-fixed infected primary hepatocytes were immunofluorescence-stained for UIS4 (this paper) and GBP1 (Clone 1B1 Santa Cruz).The samples were then treated with 0.7% Formaldehyde (v/v) (Sigma- Aldrich)/ 1% Acrylamide (v/v) (Sigma-Aldrich) in PBS for 90 minutes at 37°C. Coverslips were then incubated with a monomeric solution (19% w/w sodium acrylate (Sigma- Aldrich), 10% acrylamide (Sigma-Aldrich), 2% N, N-’-methyllenebisacrylamide (Sigma- Aldrich) in PBS), 0.5% TEMED, (VWR) and 0.5% ammonium persulfate (APS)(Sigma- Aldrich) for 30 minutes at 37°C. Then, coverslips were transferred into the wells of a 6- well plate filled with the denaturation buffer (200 mM sodium dodecyl sulfate (SDS)(Sigma-Aldrich), 200 mM NaCl (Sigma-Aldrich), 50 mM Tris (pH 9, VWR) for 15 min at room temperature, in agitation. The gels were then separated from coverslips and transferred into micro tubes containing the denaturation buffer and denatured at 72 °C for 30 min. The denatured gels were transferred into Petri dishes with 20 mL MilliQ water shaking for 30 min, 3 times. The expanded gels were shrunk by washing thrice in 20 mL of PBS for 15 min. The gels were blocked with 2% BSA-PBS for 1 h at room temperature on an orbital shaker. After blocking, primary antibodies (1:500 anti-UIS4 (this paper) plus 1:100 anti-GBP1(Clone G12 Santa Cruz) in 1% BSA-PBS blocking buffer were added and the gels were incubated for 180 min at 37°C on an orbital shaker. The gels were washed thrice in 0.5% PBS-Tween 20 (PBS-T), for 10 min, then incubated with secondary antibodies diluted in 2% BSA/PBS. The gels were then transferred into Petri dishes filled with 20 mL PBS and placed on a platform shaker for 5 min, repeated 3 times. Gels were transferred to microtubes and stained with 5µg/mL DAPI (Sigma-Aldrich) diluted in 2% BSA (Fisher Scientific)/PBS at 37°C on an orbital shaker for 60 min, transferred to Petri dishes with 20mL milliQ water, and washed for 30 minutes 3 times. To prepare gels for imaging, small sections were cut from the larger gel and dried before being placed into poly-d-lysine precoated 35 mm #1.5 coverslip bottomed dishes. Images were acquired on a Zeiss LSM980 microscope with an Airyscan detector. Image analysis was performed using ZEN Blue (Version 3.1) software.

### ROS production

ROS production was assessed by CellROX Green (InVitrogen) staining following the manufacturer’s protocol. In short, primary murine hepatocytes were plated in collagen-coated black-walled 96 well plates for 24h. Following *Pb*-infection or not, the cultures were treated with vehicle, specific inhibitors from 6h p.i (or as specified), or type-I IFNs (PBL) and each well was incubated for 30 minutes with CellROX at 37°C, 5% CO2, washed thrice with PBS and fluorescence (Ex/Em 485/520) measured using the Spectramax iD3 (Molecular Devices) and the SoftMax Pro 7.1.2 software. Tert-Butyl hydroperoxide (TBT) (Acros Organics) was used as a positive control to induce oxidative stress.

### Transfection of cells

Primary hepatocytes (7×10^5^ cells) were transfected with various plasmids (at 6 μM concentration) and siRNA (Dharmacon siGENOMESMARTpool) using the Mouse/ Rat hepatocyte Nucleofector kit (Lonza) following the manufacturer’s protocol. The transfected cells were transferred to collagen-coated wells and allowed to recover for 24 hr prior to infection or treatments.

### Therapeutic regimens

The following treatment regimens were used in this study: GBP1, LC3, Galectin-3 or scrambled siRNA (Dharmacon siGENOMESMARTpool), 1nM/mouse, hydrodynamic i.v., -1 or −2d p.i. 100mM NAC (Sigma-Aldrich) and 100mM NH4Cl (Sigma-Aldrich) in the drinking water −24hpi up until 36 hpi. 0.3mg/Kg Bafilomycin A1 (ChemCruz): 0, 12, and 24h p.i. (intraperitoneally, i.p),10mg/Kg GLX35122 (MedChem Express): 0h p.i. (i.p.), 100mg/Kg GSK2795039 (Tocris): 0h. p.i (i.p.), 20mM NAC 6 and 12h p.i (i.p.), NH4Cl 20mM at 6 and 12h p.i. (i.p.), murine IFN-I 3.3μg/Kg 0,12 and 24h p.i (i.p.).

### Hydrodynamic delivery of nucleic acids

Hydrodynamic injections were performed as described in detail before (Kim and Ahituv, 2013; Marques-da-Silva *et al*., 2024a; Sebestyen et al., 2006). In short, the desired amount of the siRNA was resuspended in PBS (at 10% volume/ body weight of the mouse) and delivered to the tail vein of mice, using constant pressure, within 7 seconds.

The mice were then placed on a warm heating pad and allowed to recover for about 30 mins, before being transferred back into their cages.

### ELISA

ELISA was performed as described in detail before (Kurup *et al*., 2017). In short, serial dilutions of supernatants or whole cell lysates (normalized for total protein) were coated in triplicate in 96-well format in 0.1M carbonate bicarbonate buffer (pH 9.6) overnight at 4°C, washed thrice with 0.05% Tween-20 (Sigma-Aldrich) in PBS (PBS-T), blocked for 60 min with 1% BSA/PBS, probed with 1mg/mL polyclonal GBP1, GBP-2 and GBP-5 (Invitrogen), monoclonal anti-mouse α- interferon (PBL), and 1mg/mL anti-mouse LC3 (Clone 4E+12, MBL international) for 60 min at 37°C, washed 5 times with PBS-T, probed with corresponding HRP conjugated secondary antibodies (Jackson and Santa Cruz) in 1% BSA/PBS and again washed 3 times with PBS-T before developing. All ELISA assays were developed using TMB liquid substrate system (Sigma-Aldrich), stopped with 2N sulfuric acid, and then read at 450nm in Spectramax iD3 (Molecular Devices) and the SoftMax Pro 7.1.2 software

### Western blot assay

Western blots were performed as detailed before (Gurung et al., 2015). In short, cells were lysed in RIPA Lysis Buffer (10mM Tris-HCl, 1mM EDTA, 0.5mM EGTA, 1% Triton X- 100, Sodium Deoxycholate, 0.1% SDS, 140mM NaCl diluted in water) containing protease inhibitors (Roche). Lysates were run on 12% SDS-polyacrylamide gel (BioRad) by electrophoresis and transferred to Nitrocellulose (BioRad) membranes. After blocking with Intercept blocking Buffer (Licor) for 1h at room temperature, the membranes were probed with the primary antibodies: anti-mouse Caspase-1p20 (Clone Casper-1, Adipogen), mGasdermin D (Clone EPR19828, AbCam), anti-mouse LC3 (Clone 4E12, MBL international), anti-mouse NOX2 (Clone 53, BD Biosciences), anti-mouse NOX4 (Clone C-3, Santa Cruz), anti-mouse GBP1 (Clone G-12, Santa Cruz), anti-mouse Rubicon (polyclonal, Abcam) and anti-mouse β-tubulin (Clone AMC0498, ABclonal) at 4^0^C overnight, washed with tris-buffered saline containing 0.1% Tween-20 (TBST) thrice, and incubated at room temperature for 60 min with relevant correspondent secondary polyclonal IRDye (Licor) conjugated antibodies. After 3 washes with TBST, the proteins were visualized directly by fluorescence (Licor). Western Blot densitometry was performed using ImageJ software. The background signal around the band was included in the data analysis.

Immunoprecipitation was carried out as described before (Marques-da-Silva *et al*., 2023). Briefly, the samples along with 10 µg/mL of the mCaspase-1p20 (Clone Casper-1, Adipogen) antibody and protein-G-Dynabeads (Invitrogen) were incubated overnight at 4° C. The cells lysates were immunoprecipitated overnight at 4° C, after which they were washed thrice with PBS containing 0.1% Tween-20 (PBST). Bound protein was eluted by the addition of 2X SDS-sample buffer, and boiled for 5 min, and then analyzed by Western Blot analysis as described above, stained with anti-GBP1 (Polyclonal, Proteintech).

### Macrophage Differentiation

BMDMs were prepared as described previously (Gurung *et al*., 2015). In short, bone marrow cells were grown in L-cell-conditioned IMDM medium (ThermoFisher) supplemented with 10% FCS (Sigma-Aldrich), 1% nonessential amino acids and 1% penicillin–streptomycin (Gibco) for 5 d to differentiate into macrophages. On day 5, BMDMs were seeded in 6-well cell culture plates. The next day BMDMs were stimulated with 100 ng/mL LPS (Invivogen) (3.5 h) followed by 5 mM ATP (Sigma-Aldrich) and the whole-cell lysates were collected as indicated.

### Cell-death assay

Lactate dehydrogenase (LDH) release assay to determine cell lysis: Overnight cultures of 8x10^3^ hepatocytes/ well in 96-well plates were inoculated with 8x10^3^ *P. bergheii* sporozoites. Cell culture supernatants were replaced with fresh media at 6h of incubation along with different treatments after washing the cells twice with media, to remove free sporozoites. Culture supernatant was subsequently collected at 24 hpi and assayed for LDH as a measure of total cytolysis, using the CytoTox 96 Non-Radioactive Cytotoxicity Assay (Promega) according to the manufacturer’s instructions. %Cytolysis calculated as 100 X (experimental LDH release signal/ maximum LDH release signal).

## Statistical analyses

Data were analyzed using Prism7 software (GraphPad) and as indicated in figure legends.

---

**Figure S1.**
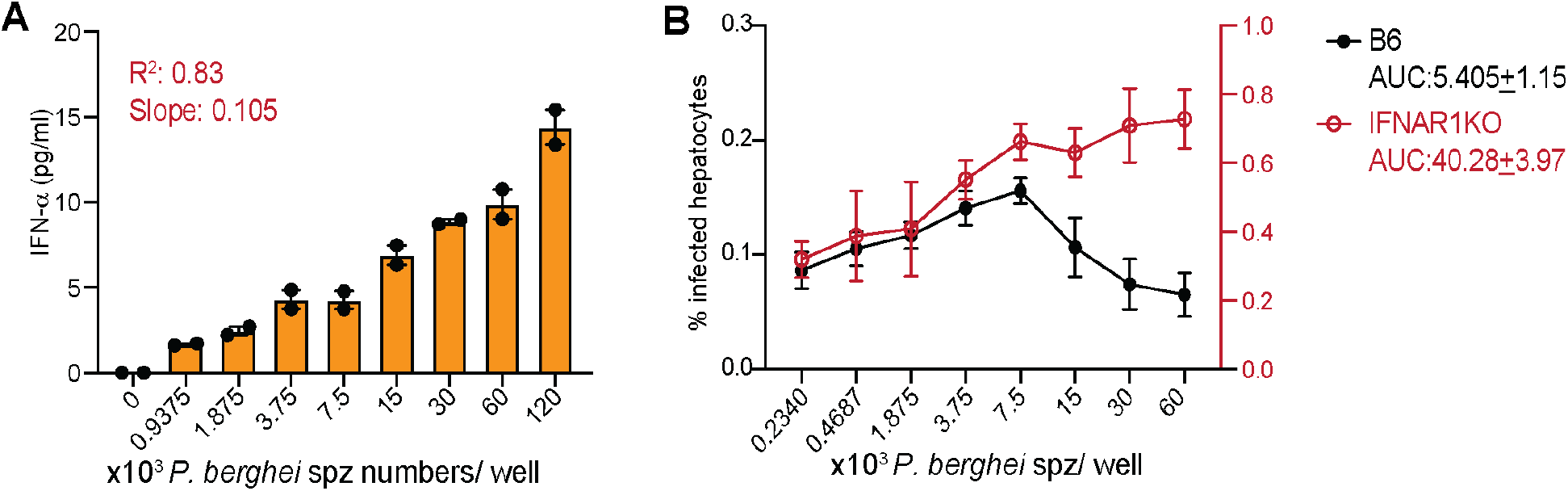
IFN-I produced by infected hepatocytes impede *Pb* infection. (A) IFN-α in the supernatants of primary B6 hepatocyte culture inoculated with increasing doses of *Pb* Spz, quantified by ELISA, 18h p.i. Each well plated with 1.2 x 10^4^ hepatocytes. Combined data presented as mean + s.e.m, from 2 separate replicate experiments, analyzed for regression with non-linear fit yielding the indicated R2 and slope values. (B) Relative parasite burdens in primary B6 or IFNARKO hepatocyte cultures inoculated with *Pb-Luc* at the indicated doses, quantified by immunofluorescence signal at 36h p.i. Each well plated with 1.2 x 10^4^ hepatocytes. Combined data from two replicate experiments presented as mean + s.e.m. AUC: area under the curve + s.e.m.

**Figure S2.**
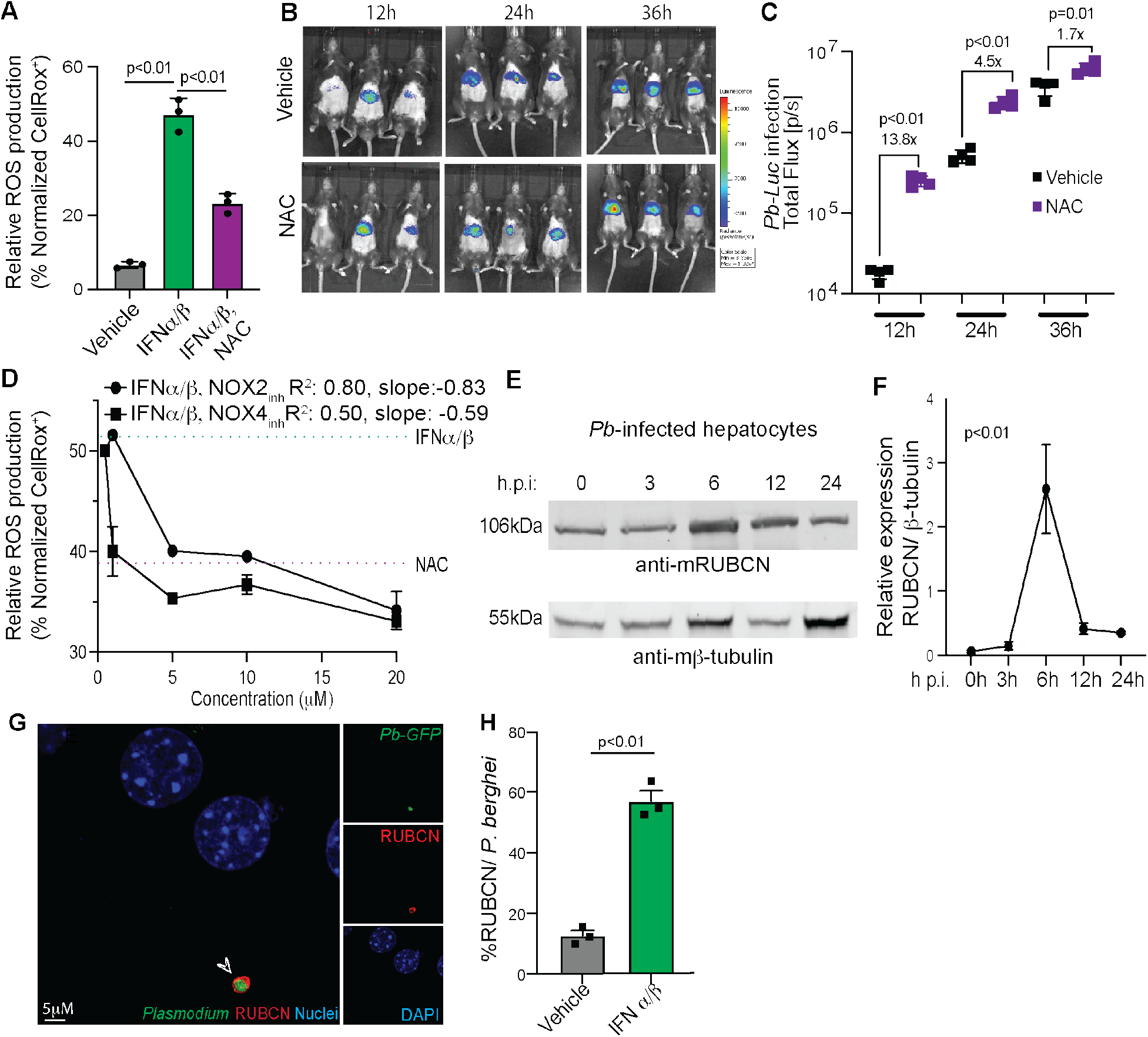
ROS, RUBCN, and NOX facilitate IFN-I-mediated control of *Plasmodium* in hepatocytes. (A) Relative ROS production in *Pb*-infected primary murine hepatocyte cultures, 24h p.i., following treatment with vehicle or IFNα/β at 6h p.i. Combined data presented as mean + s.e.m with each point representing a separate experiment, analyzed using 1-way ANOVA with Tukey’s correction to obtain the indicated p-values. (B) Representative rainbow images of luminescence signal representing liver parasite burdens at the indicated time points in (B6) mice treated with NAC (from −24h p.i.) and inoculated with *Pb-luc.* Data representative of 3 separate experiments. (C) Scatter plots showing relative liver parasite burdens at the indicated time points in (B). Representative data shown as mean + SD from 1 or 3 experiments with each data point representing a mouse, analyzed using 1-way ANOVA with Tukey’s correction to obtain the indicated p-values. (D) Relative ROS production in *Pb*-infected primary mouse (B6) hepatocyte cultures at 24h p.i., following treatment with the indicated concentrations of NOX inhibitors. Representative data from 1 of 2 replicate experiments presented as mean + s.e.m, analyzed for regression with non-linear fit yielding the indicated R^2^ and slope values. (E-F) Immunoblots indicating RUBCN expression in primary mouse (B6) hepatocyte cultures inoculated with *Pb* Spz at the indicated timepoints (D). Expression of RUBCN in the above cultures determined using relative band densities with β-tubulin serving as loading controls (E). Data represent 3 replicate experiments, analyzed using ANOVA with Geisser-Greenhouse correction to yield the indicated p-value. (H) Bar graph indicating the frequency of RUBCN colocalization with *Pb-GFP* at 24h p.i. following treatment with vehicle or IFNα/β for 90 min. Data quantified from three replicate experiments, each data point represents a replicate experiment with >50 infected hepatocytes analyzed, presented as mean + s.e.m, and analyzed using unpaired t-test to obtain the indicated p-values.

**Figure S3.**
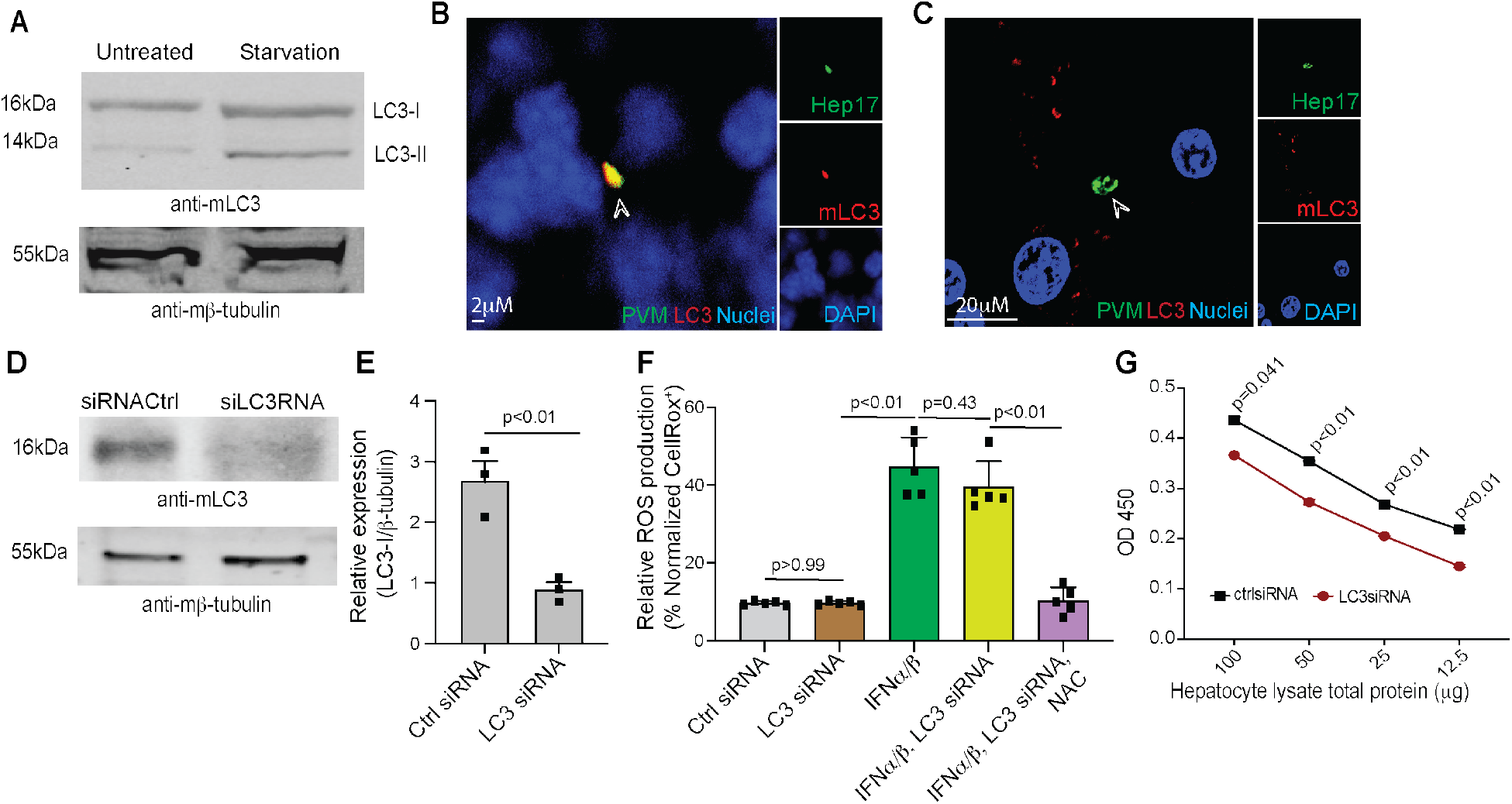
LC3 siRNA downregulates LC3 expression in hepatocytes *in vitro* and *in vivo*. (A) Immunoblot indicating LC3-I and LC3-II levels in control or starved (8h) BMDM cultures. β-tubulin served as loading controls. (B-C) Pseudo-colored confocal images depicting LC3 and *Pb* PVM (Hep17) in primary B6 (B) or IFNARKO (C) hepatocytes infected with *Pb-GFP*, 12h p.i. The arrow points to the parasite, representative image from 1 of 3 replicate experiments. (D) Immunoblot indicating LC3-I expression in primary B6 hepatocyte cultures transfected with ctrl or LC3 siRNA at 24h. β-tubulin served as loading controls. (E) LC3-I expression levels in (D) determined using relative band densities using β-tubulin as loading controls. Data represent 2 replicate experiments analyzed using student’s t- test to yield the indicated p-value. (F) ROS production in *Pb*-infected B6 primary hepatocyte cultures determined at 24 h p.i., following the indicated treatments. Combined data presented as mean + s.e.m with each point representing a biological replicate, analyzed using 2-way ANOVA with Tukey’s correction to obtain the indicated p-values. (G) Relative levels of LC3 determined by ELISA in the primary hepatocyte lysates derived from B6 mice following the delivery of the indicated siRNAs (at −48h) and LPS + poly l:C (at −24h). Representative data shown from 1 or 3 experiments with 3 mice/group, compared with 2-tailed t-tests at each dilution to yield the indicated p-values.

**Figure S4:**
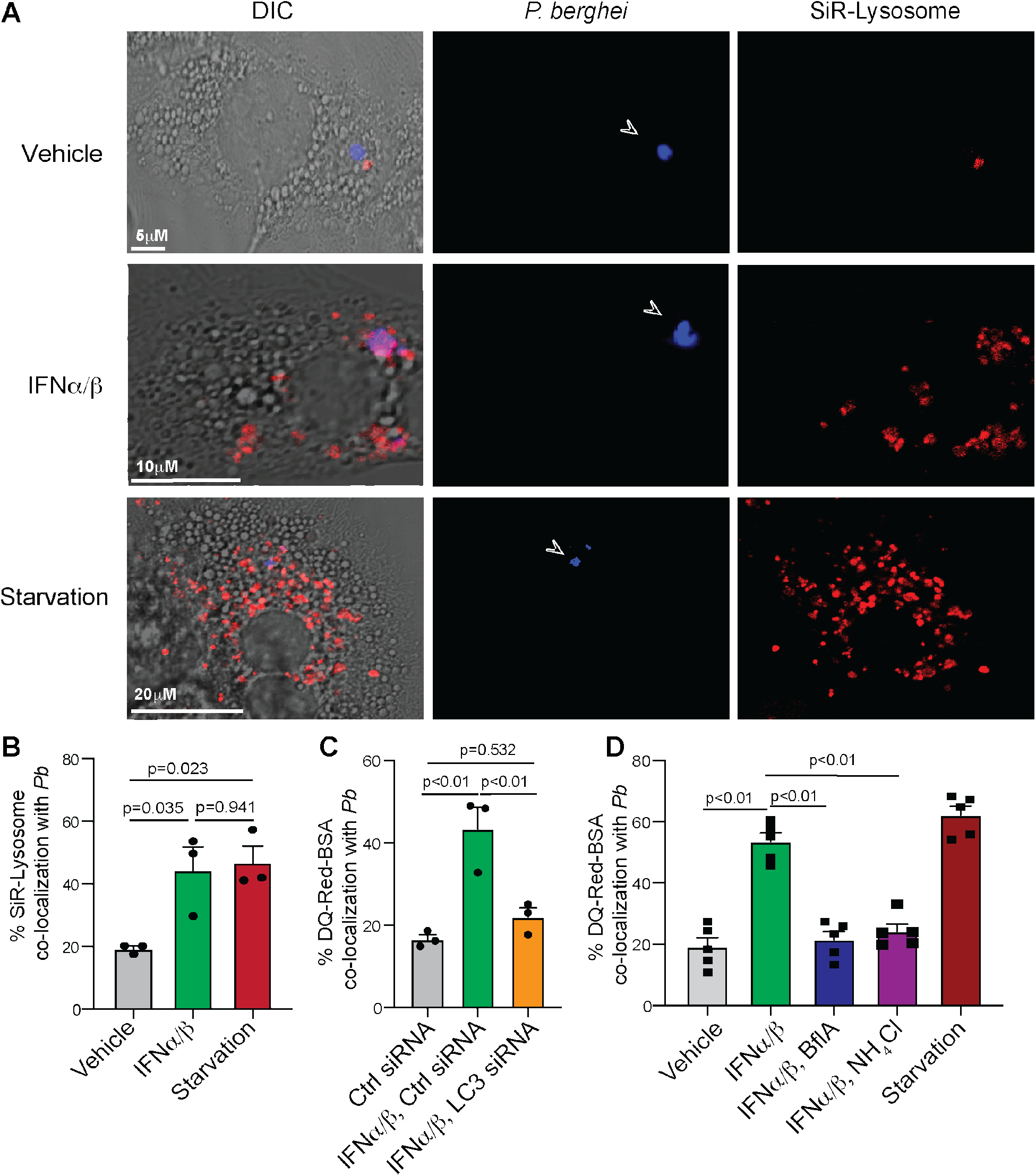
IFN-I induced lysosomal association with *Plasmodium* in hepatocytes. (A) Representative (n >10) pseudo-colored confocal images depicting live primary mouse (B6) hepatocytes co-incubated with *Pb* (CTV^+^) and IFNα/β or under starvation (24h) treated with SiR-Lysosome probe, at 24h p.i. The arrow indicates *Pb*. (B) Bar graph indicating the frequencies of SiR-lysosome colocalization with *Pb* in (A). Combined data presented from three replicate experiments with each data point depicting a replicate experiment with >50 infected hepatocytes evaluated, presented as mean+s.e.m, and analyzed using 1-way ANOVA with Tukey’s correction to obtain the indicated p-values. (C-D) Bar graph depicting the frequencies of DQ-Red colocalization with *Pb* (CTV^+^) at 24h p.i. in primary B6 mouse hepatocytes treated as indicated from 6h p.i., or starved for 12h. Combined data shown with each point representing a replicate experiment with >50 infected hepatocytes, presented as mean + s.e.m, and analyzed using 1-way ANOVA with Tukey’s correction to obtain the indicated p-values.

**Figure S5:**
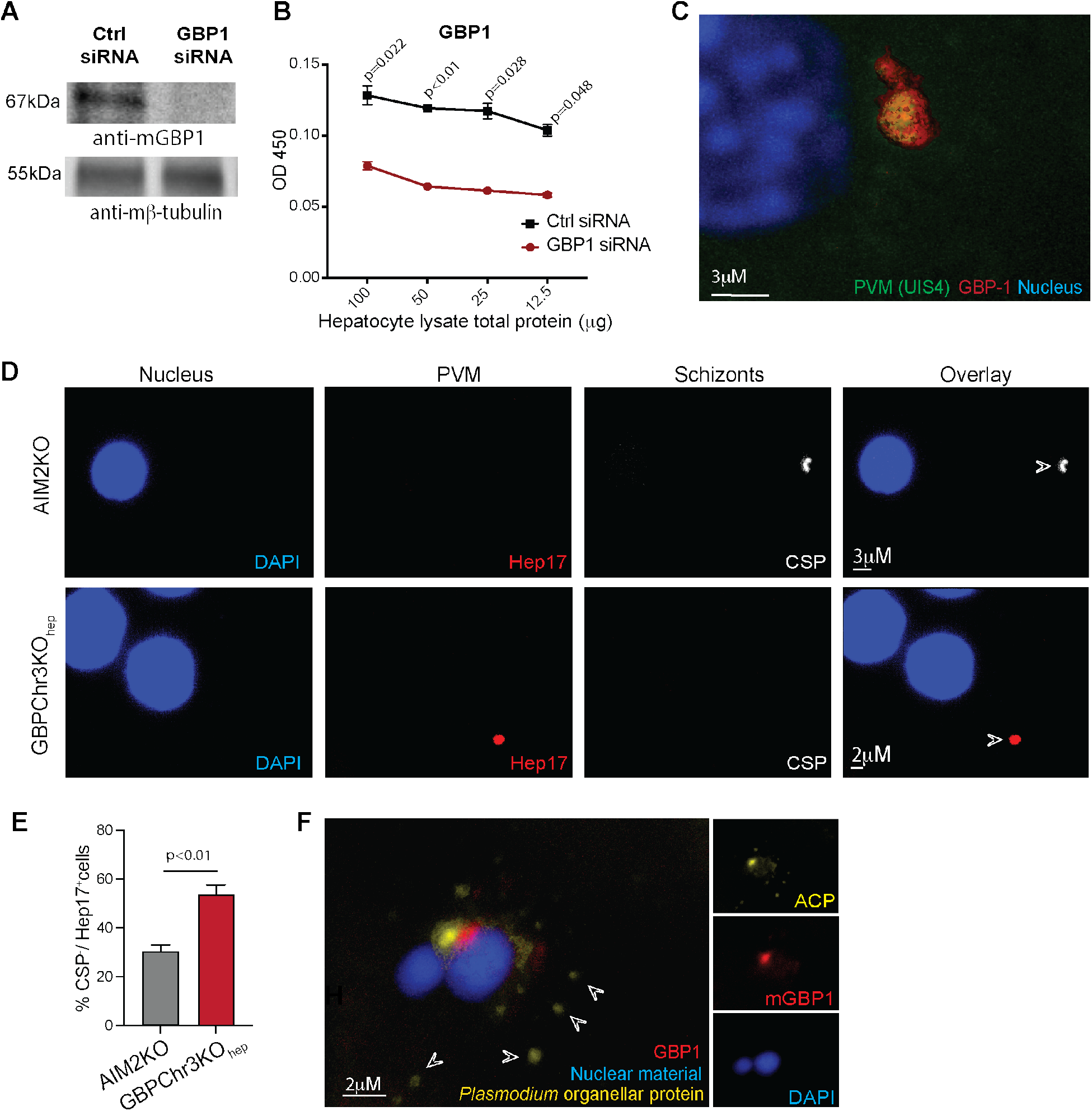
GBP1, 2, 5 localization and function in *Plasmodium* infected hepatocytes. (A) Immunoblot indicating GBP1 expression in primary B6 mouse hepatocyte cultures transfected with ctrl or GBP1 siRNA, 24h post transfection. β-tubulin served as loading control. (B) Relative protein levels of GBP1 determined by ELISA in the primary hepatocyte lysates derived from B6 mice following the delivery of the indicated siRNAs at −48h and IFNα/β at −24h. Representative data shown from 1 or 3 experiments with 3 mice/group, compared with 2-tailed t-tests at each protein concentration to yield the indicated p- values. (C) Pseudo-colored confocal image with surface rendering depicting GBP1 localization to the PVM (UIS4) of *Py* in primary B6 hepatocytes, 30h p.i. Image representative of >3 replicate experiments. (D) Pseudo-colored confocal images from 0.01% digitonin-permeabilized cultured primary hepatocyte from the indicated mice infected with *Pb*, 30hpi. Hep17 indicates intact PVM, CSP indicates *Pb* schizont surface in infected hepatocytes. Data represent > 5 separate observations. (E) Bar graphs representing the frequencies Hep17^+^*Pb* EEFs devoid of CSP staining. EEFs were considered Hep17^+^ if >30% of the circumference of the parasite was surrounded by it. Representative data shown as mean + s.e.m from 3 separate experiments with >50 infected cells (CSP^+^ or Hep17^+^) assessed, compared using 2-tailed t-test to yield the indicated p-value. (F) High-resolution confocal images depicting GBP1 localization relative to *Pb* in infected primary mouse (B6) hepatocytes in culture, 24h p.i. The arrows indicate fragments of the *Pb* organellar component, Acyl Carrier Protein (ACP).

**Figure S6:**
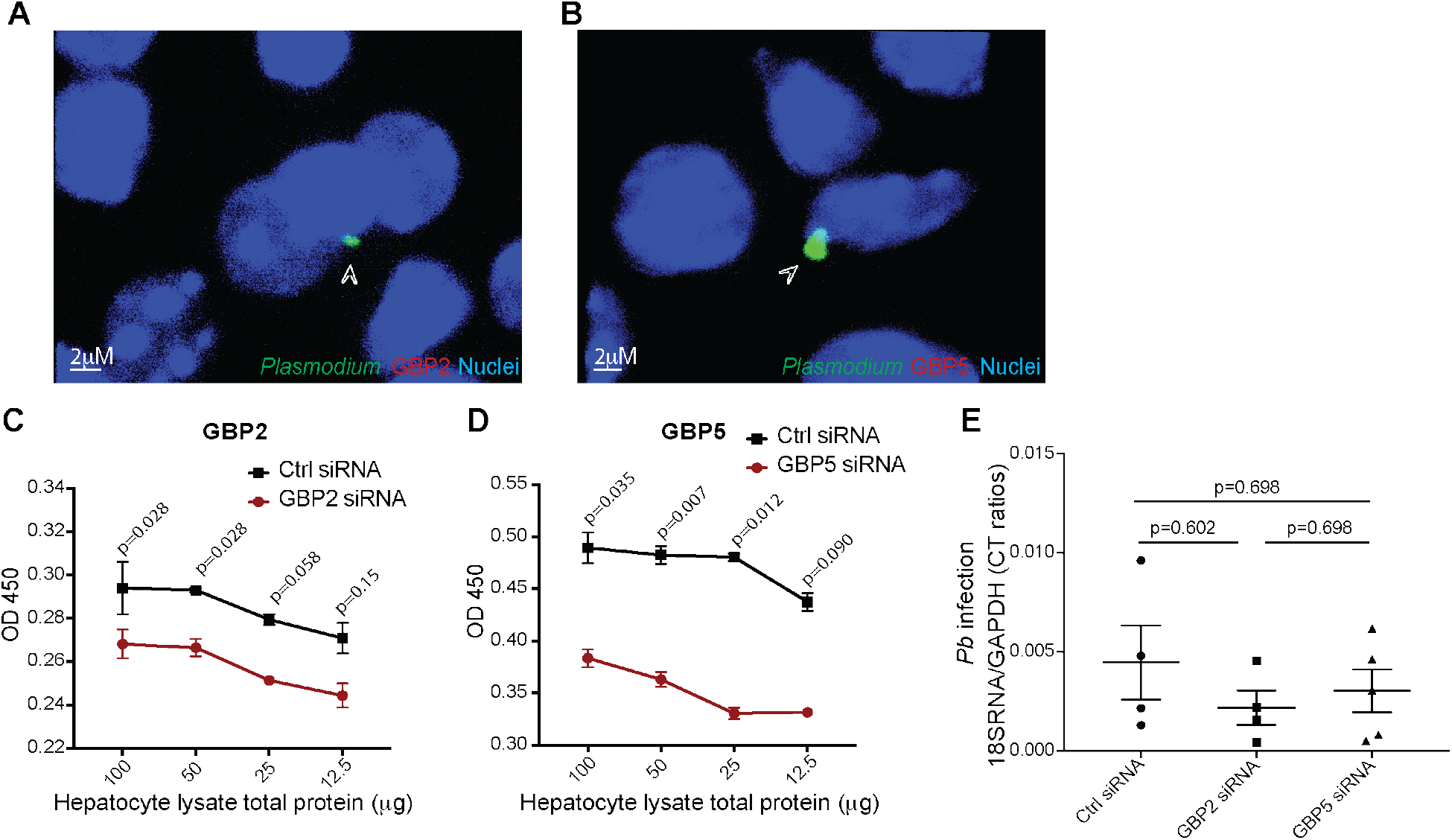
GBP2 and GBP5 localization and function in Plasmodium-infected hepatocytes. (A, B) Pseudo-colored confocal images depicting GBPs 2 (A) or 5 (B) staining in *Pb*- infected whole livers of B6 mice inoculated with *Pb-GFP*, 30h p.i. The arrow points to the parasite, images shown from 1 of >3 biological replicate experiments. (C, D) Relative protein levels of GBPs 2 (C) or 5 (D) determined by ELISA from primary hepatocyte lysates of B6 mice following the delivery of the indicated siRNAs at −48h and IFNα/β at −24h. Representative data shown from 1 or 3 experiments with 3 mice/ group, compared with 2-tailed t-tests at each protein concentration to yield the indicated p-values (E) Scatter plot indicating relative liver parasite burdens in *Pb* infected B6 mice that received control, GBP2 or GBP5 siRNAs at −48h p.i., determined by qRT-PCR at 40h p.i. Representative data from two replicate experiments with each data point representing a mouse, presented as mean + SD and analyzed using 1-way ANOVA with Tukey’s correction to obtain the indicated p-values.

**Figure S7:**
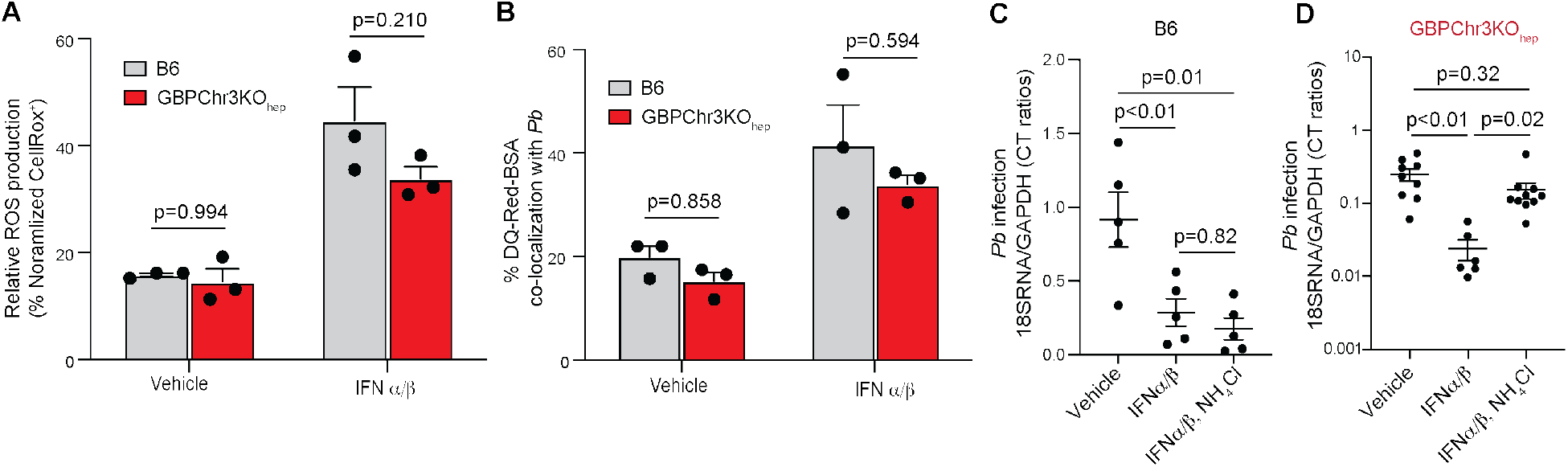
IFN-I driven ROS and lysosome-mediated control of liver-stage malaria occurs independently of GBPs. (A) ROS production in *Pb*-infected B6 or GBPChr3KOhep primary mouse hepatocyte cultures determined at 24h p.i., following treatment with vehicle or IFNα/β at 6h p.i. Combined data presented as mean + s.e.m, each point represents a separate experiment, analyzed using 2-way ANOVA with Tukey’s correction to obtain the p-values. (B) Frequency of DQ-RED colocalization with *Pb* (CTV) at 24h p.i. in B6 or GBPChr3KOhep primary mouse hepatocytes treated with vehicle or IFNα/β at 6h p.i. Combined data presented as mean + s.e.m, each point represents a replicate experiment, analyzed using 2-way ANOVA with Tukey’s correction to obtain the indicated p-values. (C-D) Scatter plot indicating relative liver parasite burdens at 40h p.i. in *Pb-*infected B6 (C) or GBPChr3KOhep (D) mice following treatment with the indicated reagents at −24h p.i. Combined data from 2 replicate experiments presented as mean + s.e.m, and analyzed using ANOVA with Tukey’s correction to obtain the indicated p-values.

**Figure S8:**
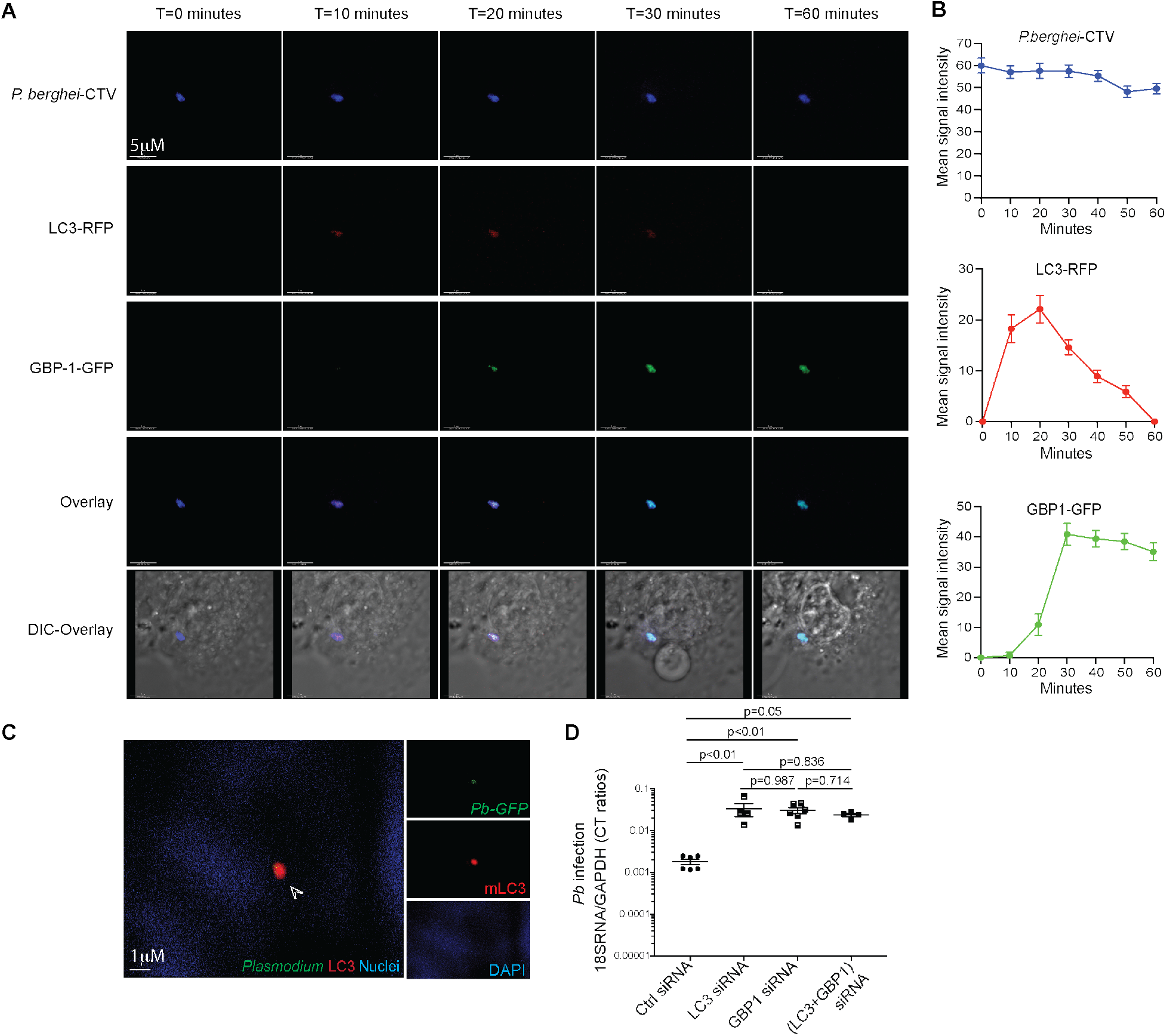
LC3 precedes GBP1 localization to *Plasmodium* and exhibits functional redundancy in limiting liver-stage malaria. (A) Representative pseudo-colored confocal time-lapse images depicting the kinetics of LC3 (RFP) or GBP1 (GFP) localization to *Pb* (CTV^+^) in infected Hepa1-6LC3-GBP1 reporter hepatocytes at the indicated time frames. Data representative of >5 infected cells from 2 separate experiments. (B) Quantification of mean fluorescence signal intensity of *Pb* (blue), LC3-RFP (red) or GBP1 GFP (green) from (A) at the indicated time frames. (C) Pseudo-colored confocal images depicting subcellular localization of LC3 and *Pb- GFP* in the liver cryosections of GBPChr3hep mice inoculated with *Pb-GFP*, 30h p.i. The arrow points to the parasite, images shown from 1 of >3 biological replicate experiments. (D) Scatter plot indicating relative liver parasite burdens at 24h p.i. in *Pb-*infected B6 mice that received control or the indicated siRNAs by hydrodynamic injection at −48h p.i. Combined data from 2 replicate experiments presented as mean + s.e.m, and analyzed using ANOVA with Tukey’s correction to obtain the indicated p-values.

**Figure S9:**
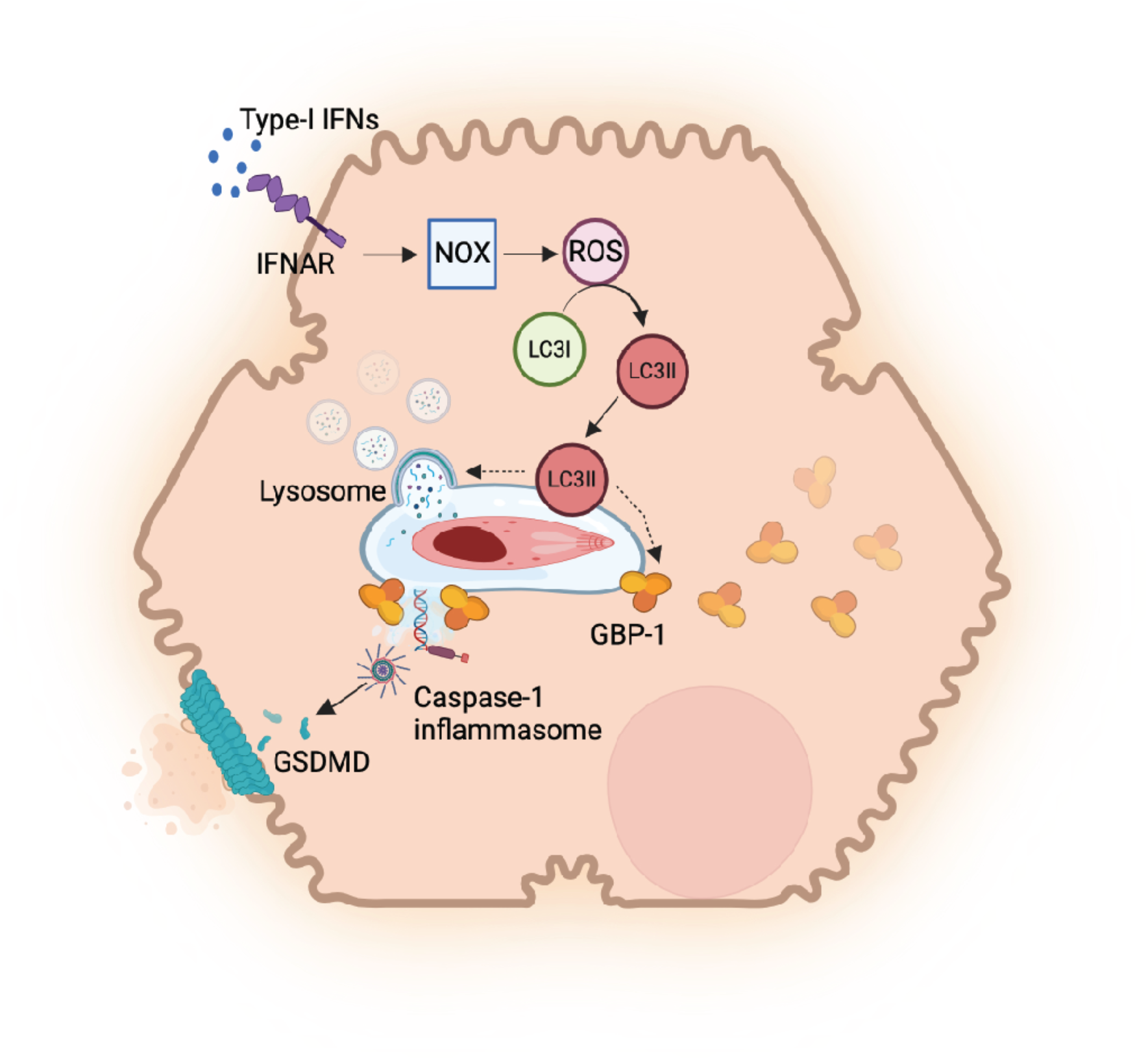
Proposed model of type-I IFN-induced control of *Plasmodium* in hepatocytes. We propose that type-I IFN stimulation in *Plasmodium*-infected hepatocytes induce NOX2 and NOX4-mediated production of ROS that activates LC3-I to LC3-II, allowing the latter to decorate the PVM. This facilitates lysosomal fusion with the PV, leading to the lysis of the parasite. Parallelly, LC3-II also promotes GBP1 association with the PVM, which disrupts the PV, potentially exposing *Plasmodium* PAMPs including its dsDNA to the hepatocyte cytosol. This triggers AIM2-mediated caspase-1 inflammasome activation and the resultant GSDMD-mediated pyroptosis in the host hepatocytes.

